# SOX7 and ERG reprograms neuroectoderm into microenvironmentally attuned brain endothelial cells that possess forebrain regional bias

**DOI:** 10.64898/2026.11.09.764683

**Authors:** Tyler M. Lu, Ugur Akcan, Alexander J. Rhee, Aomeng Cui, Jiuhai Wang, Sean Houghton, Fuqiang Geng, Victor A. Martinez, Sanjid Shahriar, Jenny Xiang, Zev Rosenwaks, Shahin Rafii, Irene M. Pedersen, James Gorman, Raj Wadgaonkar, Frank C. Barone, Peter Galie, Dritan Agalliu, Raphael Lis

**Affiliations:** Ronald O. Perelman and Claudia Cohen Center for Reproductive Medicine, Weill Cornell Medicine, New York, NY 10065; Molecular and Cellular Biology Program, SUNY Downstate School of Graduate Studies, Brooklyn, New York 11203; Department of Neurology, Columbia University Irving Medical Center, New York, NY 10032; Division of Regenerative Medicine, Department of Medicine, Weill Cornell Medicine, New York, NY 10065; Scintillon Aging Research Center, Scintillon Institute, San Diego, CA 92121; Wyss Institute for Biologically Inspired Engineering, Harvard University, Boston, MA 02115; Genomics Resources Core Facility, Weill Cornell Medicine, New York, New York 10065; Department of Biomedical Engineering, Rowan University, Glassboro, NJ, USA; Department of Pathology & Cell Biology, Columbia University Irving Medical Center, New York, NY 10032

**Keywords:** human PSC, blood-brain barrier, BMEC, neurovascular unit, endothelial identity, epithelial identity

## Abstract

Reliable human blood brain barrier (BBB) models are limited by a persistent problem of cellular identity: many protocols select cells for barrier or transporter properties rather than authentic, regionally specified brain endothelium. Here we show that constitutive expression of the transcription factors SOX7 and ERG converts human pluripotent stem cell–derived epithelial barrier-forming cells into bona-fide vascular endothelial cells (SE-rECs) whose transcriptome and proteome converge on primary human brain endothelium. Benchmarked against a human adult-brain endothelial atlas, SE-rECs acquire a forebrain-biased regional program stronger than that of permissively (ETS-)reprogrammed or umbilical-vein endothelial cells, reproducibly across biological replicates, although it does not reach the absolute regional identity of in-vivo brain endothelium. At the transcript level, epithelial-derived barrier forming cells (iBFCs) cells score highly on select barrier and transporters genes yet lack endothelial lineage identity, showing that transporter benchmarks alone do not establish endothelial fate. Co-culture of SE-rECs with astrocytes and pericytes in a two-dimensional neurovascular-unit model as well as a perfused microfluidic device induces mature BBB features and reduces paracellular permeability. SE-rECs thus provide a lineage-faithful, organotypically informed platform for modelling the human BBB.

## Introduction

The brain vascular endothelium was among the first vascular beds to be described in organotypic terms, effectively pioneering the notion of organ-specific vascular function^1^. Of these endothelial cells (ECs), brain microvascular endothelial cells (BMECs) have been characterized as non-fenestrated ECs endowed with organotypic properties that enable a strict control over the traffic of ions, molecules and cells between the central nervous system (CNS) and the bloodstream^2–10^. BMECs also possess a distinct developmental origin compared to most other mesoderm-derived endothelium and have been shown to lack the expression of nearly all HOX family genes, a characteristic that persists even in culture, distinguishing them from even spinal cord ECs which express posterior HOX genes^11,12^. Under inflammatory conditions, BMECs undergo dynamic molecular changes resulting in increased paracellular permeability and induction of adhesion molecules such as ICAM-1, VCAM-1, and selectins^13,14^.

The microvascular system in the brain is primarily composed of a complex network of ECs, astrocytes, and pericytes that come together to be collectively referred to as the blood-brain barrier (BBB)^15–20^. This dynamic multicellular structure relies on the molecular interplay between these cell types and acts as an elaborate discerning barrier that maintains a tight regulation of CNS homeostasis. As such, the BBB is crucial for optimal neuronal function and safeguarding the CNS from damage caused by toxins, pathogens, inflammation, injury, and diseases^19–27^. This multimodal barrier activity combines a physical barrier, characterized by the establishment of tight junctions^4,6,28–30^, a low rate of transcytosis which limits vesicle mediated transport^31–34^ and a low expression level of leukocytes adhesion molecules (LAM) that impedes the recruitment of immune cells to the CNS^35^. These properties are induced and sustained by crucial interactions with other cells such as mural cells, immune cells, glial cells, and neural cells, which collaborate within the neurovascular unit (NVU)^16,20^. The BBB also serves as a molecular barrier, with efflux transporters responsible for actively preventing the movement of certain substances that would normally pass through the membranes of brain endothelial cells by diffusion; as well as a metabolic barrier defined by the presence of enzymes that metabolize bioactive substances prior to their arrival in the brain parenchyma.

To model these highly organotypic functions *in vitro*, many groups have employed the use of human pluripotent stem cell (hPSC)-derived cells in an attempt to capture select phenotypes of interest such as tight junction mediated barrier formation or immune cell interaction. Use of these unidimensional BBB models has aided in the discovery and understanding of various aspects of BBB function. EECM-BMEC-like cells have shown the potential to be used for the study of immune cell migration across a vascular barrier such as the BBB while lacking a robust upregulation of tight junctions^36,37^. There are also models in which the establishment of a functional barrier has even superseded the need for vascular cell identity when modelling the BBB. In these cases, hPSC-derived epithelial barrier forming cells (iBFCs^38–41^) which exhibit high trans-epithelial electrical resistance (TEER) but no vascular endothelial cellular identity *in vitro* have been used to assess the permeability of small molecules and various pathophysiologies^38–60^. Altogether, these models have enabled the study of various aspects of the BBB *in vitro*. However, such a reductionist approach may overlook the highly organotypic properties inherent to ECs in the brain as well as the important contributions of other cell types such as astrocytes and pericytes or the impact of dynamic changes in BBB function under different physiological and pathological conditions^19,20,23–26,61^. A further, largely unaddressed limitation is regional: the brain vasculature is regionally specialized along the neuraxis, with molecularly distinct endothelial programs distinguishing forebrain from hindbrain territories^2,62,63^. Current hPSC-derived BBB models are neither benchmarked against nor selected for this brain-region identity, and whether a defined transcription-factor reprogramming strategy can install a forebrain endothelial program has not been established.

Recent advances in stem cell biology have shifted the emphasis from producing generic cell types toward deriving cells that recapitulate the spatiotemporal programs underlying tissue-specific identity^64^. *In vivo*, endothelial fate commitment precedes the acquisition of organ-specific programs, with pioneer ETS transcription factors (TFs) such as *ETV2* driving early vascular specification^65–68^. However, timely downregulation of *ETV2* is critical for organotypic maturation and underscores the necessity for alternative transcriptional circuits that sustain endothelial identity while enabling tissue-specific specialization^69–71^. This mechanism has been adapted to successfully generate generic ECs from hPSCs through a carefully timed directed differentiation^72,73^. Previous work has also demonstrated that other ETS TFs such as *ERG* maintain expression in adult vascular beds and occupies a central role as a master regulator of endothelial identity and vascular stability^74–77^. Utilizing an established ETS transcription factor-based cellular reprogramming strategy^78,79^, we were previously able to generate bona fide ECs from a neuroepithelial starting population by overexpression of *ETV2*, *ERG*, and *FLI1* (EEF-rEC)^80^. These EEF-rECs were transcriptomically and functionally vascular endothelium but lacked many aspects of brain organotypicity including a robust barrier phenotype.

SOX family TFs are well-established regulators of cell fate decisions during development^81^ with a group of them being especially involved in vascular development^82–84^. This SOXF group of TFs (*SOX7*, *SOX17*, *SOX18*) play a complementary role in vascular development and endothelial homeostasis with their combined activity shown to be essential for angiogenic patterning, arterial–venous specification, and vascular integrity with *SOX7* being especially critical in brain ECs^85–89^. These observations suggest that temporally restricted, tissue-context–dependent expression of SOXF factors provides a mechanism for endothelial fate acquisition that preserves organotypic programs, distinguishing them from the transient, generic induction mediated by pioneer factors such as *ETV2*. Use of a synergistic SOXF/ETS TF driven cellular reprogramming strategy was recently reported to generate phenotypically marked ECs from fibroblasts in the cardiac system. These cells were able to demonstrate vascular phenotypes and angiogenic potential in both *in vitro* and *in vivo* conditions while also adopting select organotypic traits^90^.

In this article, we present a novel platform for modeling the human BBB that combines the reprogramming of iBFCs into stable ECs by overexpression of SOXF/ETS TFs and Organ-on-Chip technology. This work seeks to demonstrate that reconstituting developmental SOXF/ETS circuitry *in vitro* results in ECs that not only possess an adult vascular fate but also retain a tissue-specific transcriptomic profile. Here, a synergistic SOX7, *ERG* induction strategy successfully converts neuroepithelial barrier forming cells into phenotypically marked ECs (SE-rECs) that are transcriptomically congruent to adult ECs from various human vascular beds while retaining aspects of brain-specific spatial patterning. When seeded into a microfluidics device along with the other NVU cell types (Astrocytes, Pericytes), SE-rECs adopt select brain organotypic properties, such as decreased permeability, and upregulated expression of transporters and junctional markers when compared to other ECs. In this study, we introduce a reprogramming strategy that yields spatially-aligned cells with an organotypically faithful identity as well as a BBB-Chip that can capture the multifaceted biology of the brain vasculature which could provide a more comprehensive understanding of the blood-brain barrier’s involvement in both health and disease, which can ultimately aid in the development of novel drug treatment.

## Results

To generate a hPSC-derived brain organotypic, barrier forming EC, we first incorporated bulk RNAseq data of iBFCs we generated into a dataset comprised of the global transcriptomes of all publicly available iBFC samples from a static 2D monoculture. We combined these samples with various human umbilical vein ECs (HUVECs) and adult ECs from different vascular beds and performed a bioinformatic meta-analysis to uncover the transcriptomic variances that drive the divergence between an epithelial and endothelial identity. Importantly, of the 2 *in vitro* BMEC samples included, 1 was obtained from previously published work by another group^41^ while the other was cultured by our group. The *in vivo* BMEC samples were all obtained from previously published works and included multiple patient donors^91,92^. This analysis was then focused on ETS and SOX TFs as it has been demonstrated that these families of genes regulate vascular development *in vivo* (**Fig. S1A**). *SOX7*, *SOX18*, *ERG*, *SOX17*, *ETS2*, *ETS1*, and *FLI1* were found to be highly differentially upregulated across the EC samples compared to our epithelial cell controls (**Fig. S1B, S1C**). Based on landmark studies in the field we used *SOX7* and *ERG* in a reprogramming strategy to establish a vascular EC identity in iBFCs seeking to capitalize on the functional barrier phenotype in these cells. To accomplish this, we started with the established iBFC protocol and transduced the cells with *SOX7* and *ERG* (SE) on day 8 of the differentiation while subculturing them onto a 0.1% gelatin matrix using a serum-free endothelial media under TGF-β signaling inhibition (**Fig. 1A**). At the end of this reprogramming protocol, a robust population of EPCAM^-^PECAM1^+^CDH5^+^ cells can be isolated by fluorescently activated cell sorting (FACS) and expanded into a pure population of cells expressing EC marker proteins (**Fig. 1B**). EPCAM^-^PECAM1^+^CDH5^+^ population of *SOX7* and *ERG* reprogrammed endothelial cells (SE-rECs) could be generated from both induced pluripotent stem cells (IMR90-4, H6) and embryonic stem cells (H1). SE-rECs generated from all hPSC lines maintained expression of PECAM1 and CDH5 while suppressing expression of EPCAM (**Fig. 1C**). Transduction with SOX7 and ERG together greatly improved the overall yield of phenotypically marked, putative ECs when compared to *SOX7* or *ERG* alone in the same reprogramming strategy. Transduction with *SOX7* alone led to an average yield of 20% ECs from the EPCAM^-^cells in our differentiation whereas *ERG* alone resulted in an average yield of 50% ECs in the same settings. Yields in both single-factor conditions were significantly lower than the average 75% yield we observe in our synergistic transduction approach with both *SOX7* and *ERG* (**Fig. 1D**). To assess the transcriptomic profile of these SE-rECs, we compiled high throughput RNA sequencing data including *in vitro* ECs from isolated various human tissues and *in vivo* brain ECs obtained from human biopsies. After isolation and expansion, SE-rECs generated using the IMR90-4, H6, and H1 lines were sequenced using Next Generation Bulk RNAseq technologies. These global transcriptomes were compared to all other EC samples in our dataset using a Pearson correlation matrix generated from log counts per million reads. This demonstrated that SE-rECs derived from all hPSC lines used in this study correlate with a high degree of confidence to all other endothelial cell samples based on global gene expression (**Fig. 1E**).

**Figure 1:**
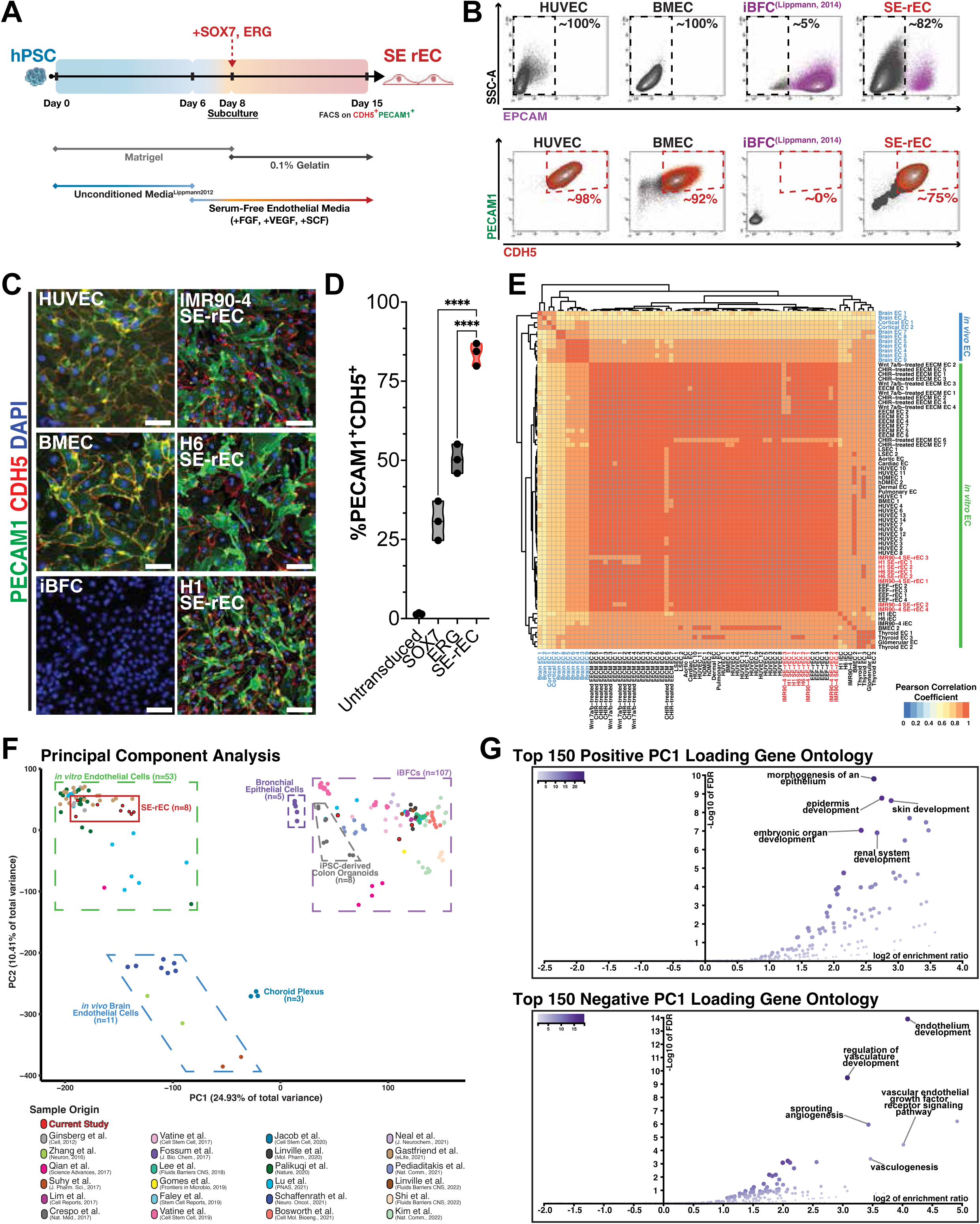
Comprehensive reprogramming of neuroepithelium using SOX7 and ERG results in transcriptomically congruent and phenotypically marked endothelial cells. **A.** Schematic of the transcription factor-mediated reprogramming method utilizing SOX7 and ERG to generate SE-rECs from iBFCs. **B.** Flow cytometry plots demonstrating expression of EPCAM, PECAM1, and CDH5 across HUVEC, BMEC, iBFC and IMR90-4 derived SE-rEC samples; N=5 biological replicates. **C.** Immunofluorescence of HUVECs, BMECs, iBFCs, SE-rECs derived from iPSCs (IMR90-4, H6) as well as ESCs (H1) with PECAM1(Green), CDH5(Red), DAPI(Blue) (scale bars=100µm); N=5 biological replicates. **D.** Violin plot showing yield of SE-rECs from single and synchronous transduction; N=3 biological replicates. ****P-value<0.0001 **E.** Pearson correlation heatmap of global transcriptomes from both in vivo and in vitro endothelial cell samples demonstrating SE-rECs derived from IMR90-4, H6, and H1 cells all highly correlate to an endothelial identity. **F.** Principal Component Analysis plot approximating relative relationship of 195 distinct cell samples across 27 library preparations from previously published and newly generated bulk RNA sequencing data. **G.** Gene-set enrichment analysis showing molecular pathways associated with the top 150 positive and negative PC1 loading genes across all samples. Transcript mapping quality parameters included a maximum number of mismatches allowed set to 10, a mismatch-to-read-length ratio of 1.0, and only uniquely mapped reads (MAPQ = 255)

To substantiate the vascular endothelial identity of the SE reprogrammed ECs (SE rECs), we again utilized a bulk RNA sequencing meta-analysis to compare our SE-rECs to *bona-fide* ECs as well as other cell types. A comprehensive dataset of global transcriptomic profiles was assembled, including one-hundred ninety-five samples from twenty-seven libraries captured by high throughput RNA sequencing. Our combined dataset included all EC samples previously used in our Pearson correlation analysis as well as one-hundred and seven iBFCs as well as other epithelial cell samples. *De novo* sample generated by our group corresponding to those from previously deposited samples were also used in this analysis which served to validate that our initial sample processing had mitigated any sequencing bias which could have impaired our downstream analyses. These samples included iBFCs differentiated from two induced-pluripotent stem cell (iPSC) lines (IMR90-4 and H6) along with various EC samples. This diverse set of libraries likely contained sequencing bias due to differences in variables such as the method of RNA preparation which we addressed by batch correction followed by a principal component analysis (PCA) to assess sample clustering in an unsupervised analysis of global transcript expression profiles (**Fig. 1F**).

In our robust meta-analysis consisting of samples generated in our lab and obtained from previously published online repositories, much of the molecular distances across all samples are captured within PC1 (24.9%), PC2 (10.4%) and PC3 (7.3%) (**Fig. S1D**). Upon mapping the top 150 most positive and negative PC1 loading genes to known molecular pathways using a gene-set enrichment analysis^93,94^, we report that SE-rECs (negative PC1 loading score) mapped to pathways associated with “endothelium development”, “VEGF receptor signaling pathway”, and “vasculogenesis” while our epithelial controls mapped to processes associated with “morphogenesis of an epithelium”, “ epidermis development”, and “skin development” (**Fig. 1G**). Further investigation into these PC1 loading genes from the PCA demonstrated a clear cellular identity divergence across the axis. This divergence was characterized by primarily endothelial (*PECAM1*, *VWF*, *SOX17*) and epithelial (*EPCAM*, *CLDN6*, *CDH1*) marker genes (**Fig. 2A**) whereas PC2 separated samples based on genes associated with cellular transport (*AQP4*) and extracellular matrix organization (*SPOCK2*) (**Fig. S1E**). A Pearson correlation analysis conducted on this combined dataset demonstrated a divergence between the highly correlative transcriptomic profiles of all endothelial samples, including SE-rECs, and all epithelial samples which also clustered with a high degree of correlation to one another (**Fig. S2**). This further confirms that SE-rECs are transcriptomically congruent with vascular ECs through another method of unbiased transcriptomic clustering. These analyses established a clear divergence between EC and epithelial cell clusters as well as some transcriptomic separation between *in vivo* isolated and *in vitro* samples. SE-rECs were shown to express canonical endothelial genes such as *CDH5*, *PECAM1*, and *CLDN5* (**Fig. 2B**) whereas all hPSC-derived iBFCs demonstrated high expression of canonical epithelial genes such as *CDH1*, *EPCAM*, and *CLDN6* (**Fig. 2C**). Our dataset included SE-rECs and iBFCs differentiated from the same starting hPSC lines indicating that these differences in expression profiles did not arise from intrinsic characteristics of the hPSCs (**Fig. S3A, B**).

**Figure 2:**
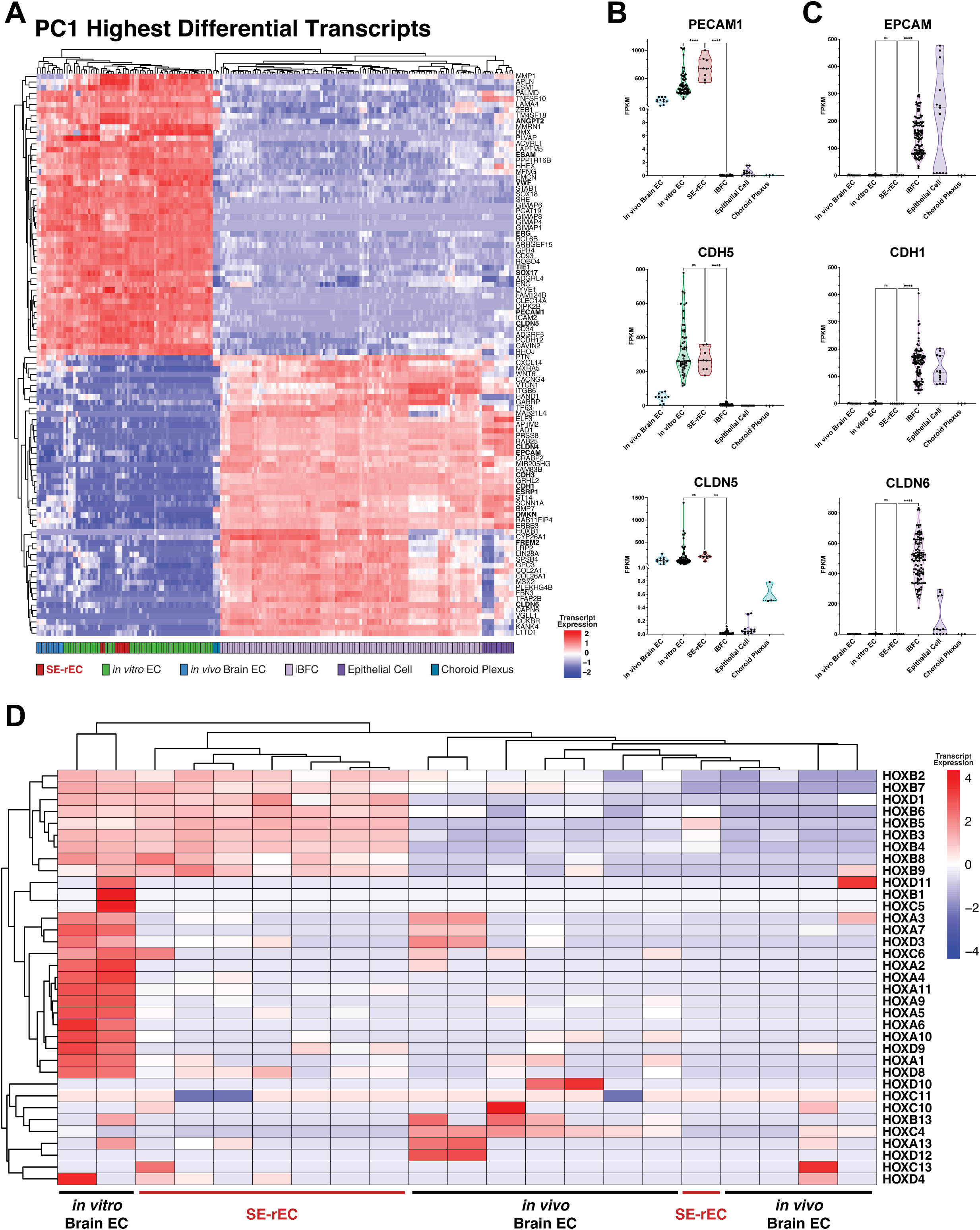
SOX7 and ERG mediated reprogramming reinforces expression of canonical endothelial transcripts while retaining spatial patterning. **A.** Heatmap of global transcriptomes from all samples in the meta-analysis demonstrating differential expression of the top 50 genes driving separation of the positive and negative ends of PC1 of the Principal Component Analysis highlighting clear separation in cellular identity between endothelial and epithelial samples. **B.** Violin plots demonstrating differential expression of endothelial marker genes highlighting significant difference between SE-rECs and iBFCs. *P-value<0.05; **<0.002; ***<0.0002; ****<0.0001. **C.** Violin plots demonstrating differential expression of epithelial marker genes highlighting significant difference between SE-rECs and iBFCs. *P-value<0.05; **<0.002; ***<0.0002; ****<0.0001. **D.** Heatmap of all HOX family transcripts mapped in the bulk RNAseq analysis demonstrating upregulated expression of various HOX genes in BMECs upon in vitro culture. SE-rECs cluster with in vivo BMEC samples by unsupervised clustering of HOX expression repertoire.

As another metric of organ specificity, we sought to investigate the HOX gene repertoire expressed by SE-rECs as compared to our organotypic endothelium of interest, brain-specific ECs. A differential gene expression analysis of the bulk RNAseq data using EdgeR demonstrated that SE-rECs express a significantly lower level of all HOX gene transcripts relative to BMECs cultured *in vitro* (FDR-adjusted p-value<0.05); however, this differential HOX gene expression was not observed when SE-rECs were compared to freshly isolated *in vivo* BMEC samples (**Fig. 2D**). All the *in vivo* BMEC samples that had never been cultured retained their developmental HOX patterning demonstrating significantly lower (FDR-adjusted p-value<0.05) or absent expression of all HOX transcripts. This generalized absence of HOX gene expression is comparable to previously reported data on cranial neural crest cells indicating that the *in vivo* BMECs used in our meta-analysis could have retained some characteristics from their embryonic origin^95^. Moreover, this would suggest that the SE-rECs have also retained some genetic traits from their embryonic origins as the iBFCs originate from neuroectoderm^96^. In an unsupervised clustering based on Euclidean distance, all SE-rEC samples clustered closer to *in vivo* BMEC samples based on HOX gene expression than either of the other *in vitro* cell types (**Fig. 2D**).

We next sought to both functionally assay our SE-rECs while also validating that the RNA transcript expression profiles correlated to functional protein expression, as with certain classes of proteins this may not always be the case^97–100^. High resolution mass spectrometry was performed on SE-rECs along with various endothelial and epithelial controls resulting in 9,713 unique protein isoforms quantified. A Pearson correlation analysis of these proteomes was conducted which revealed SE-rECs correlate to endothelial samples (BMECs, hCMEC/D3) and not to epithelial (H-1975) or hPSC (IMR90-4) samples by protein expression across all mapped proteins (**Fig. 3A**). A PCA of all mapped proteins additionally reinforces the proteomic endothelial identity of SE-rECs as they cluster with ECs along PC1 which accounts for 39.52% of the molecular distance between samples (**Fig. 3B**). The top 100 most differentially abundant proteins from PC1 as determined by loading score included endothelial markers (CDH5, ERG) and epithelial markers (EPCAM, CLDN6) reflecting the transcriptomic findings (**Fig. 3C**). SE-rECs also clustered with primary BMEC and undifferentiated hPSC along PC2 which accounted for 24.25% of the molecular distance between samples (**Fig. S4A)**.

**Figure 3:**
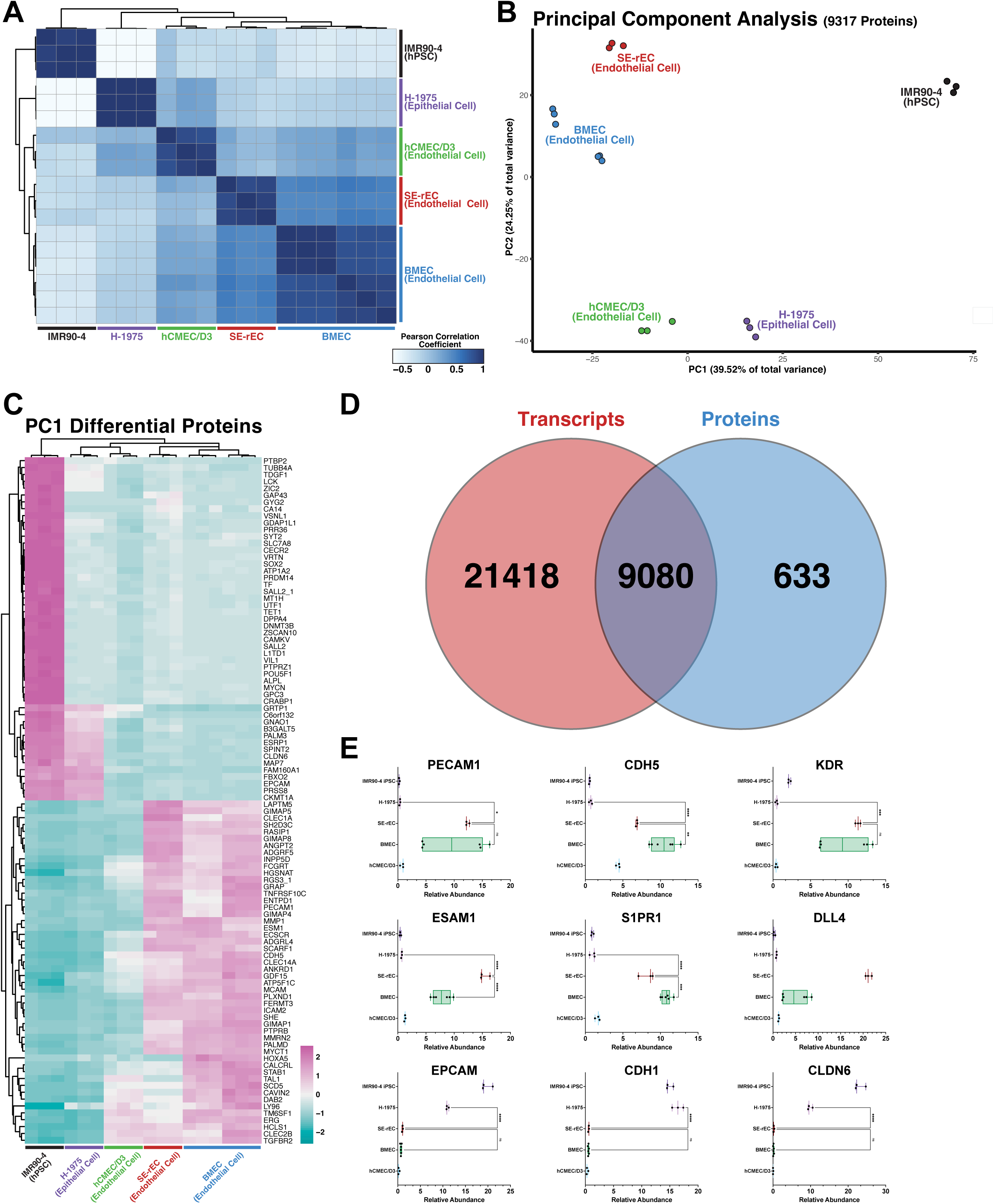
Protein abundance profiles corroborate transcriptional vascular endothelial identity of SE-rECs. **A**. Pearson correlation heatmap of proteomic profiles generated using mass spectrometry of in vitro BMECs (N=6), hCMEC/D3 (N=3), SE-rECs endothelial samples (N=3), H-1975 epithelial samples (N=3), and IMR90-4 iPSC samples (N=3). **B.** Principal Component Analysis of figure profiles demonstrating clustering of in vitro BMECs, hCMEC/D3, SE-rECs endothelial samples separately from H-1975 epithelial samples, and IMR90-4 iPSC samples across PC1. **C.** Heatmap showing differential relative abundance of the top and bottom 50 PC1 loading score proteins across BMEC, hCMEC/D3, SE-rEC, H-1975, and IMR90-4 samples. **D.** Venn diagram illustrating correspondence of mapped proteins to transcripts from the bulk RNAseq dataset. **E.** Box and whisker plots demonstrating relative abundance of select endothelial and epithelial proteins across all protein samples. *P-value<0.05; **<0.002; ***<0.0002; ****<0.0001.

Upon direct comparison, greater than 97% (9080) of all unique protein groups mapped in this analysis corresponded to the transcripts mapped from the bulk RNAseq analysis demonstrating an independent validation of both our RNA and Protein datasets (**Fig. 3D**). Proteins associated with transcytosis, as annotated by the Gene Ontology Resource, similarly clustered SE-rECs with BMECs (**Fig. S4B**). This result would be recapitulated when we evaluated the abundance of all SLC transporter proteins uniquely mapped in our dataset where we report that SE-rECs have a similar proteomic repertoire to BMECs *in vitro* (**Fig. S4C**). Relative abundance of endothelial specific proteins such as CDH5, PECAM1, and KDR were revealed to be high in SE-rECs as well as BMECs while being low in iPSCs and epithelial cells. Congruently, abundance of epithelial specific proteins such as CDH1, EPCAM, and CLDN6 was low in SE-rECs and the endothelial controls validating the results from our transcriptomics analysis (**Fig. 3E**).

Through this proteomic assay, SE-rECs were also found to possess select brain-specific transporters and tight junction-associated proteins such as SLC2A1, CLDN5, and OCLN (**Fig. 4A**). We were then able to confirm expression of CLDN5 and OCLN localized to the cell-cell junctions in SE-rECs compared to primary human BMECs using confocal microscopy (**Fig. 4B**). Furthermore, these hPSC-derived cells were able to form stable vessels on Matrigel *in vitro* indicating angiogenic potential even under serum free conditions (**Fig. 4C**). SE-rECs recapitulated a vascular-specific response to inflammatory stimulation demonstrated by upregulation of E-selectin to the cellular membrane after acute stimulation with 100ng.mL^-1^ of TNF-α (**Fig. 4D, 4E**). The presence of these proteins indicates a robust endothelial identity in hPSC-derived SE-rECs generated from our reprogramming strategy that have been expanded and maintained *in vitro*.

**Figure 4:**
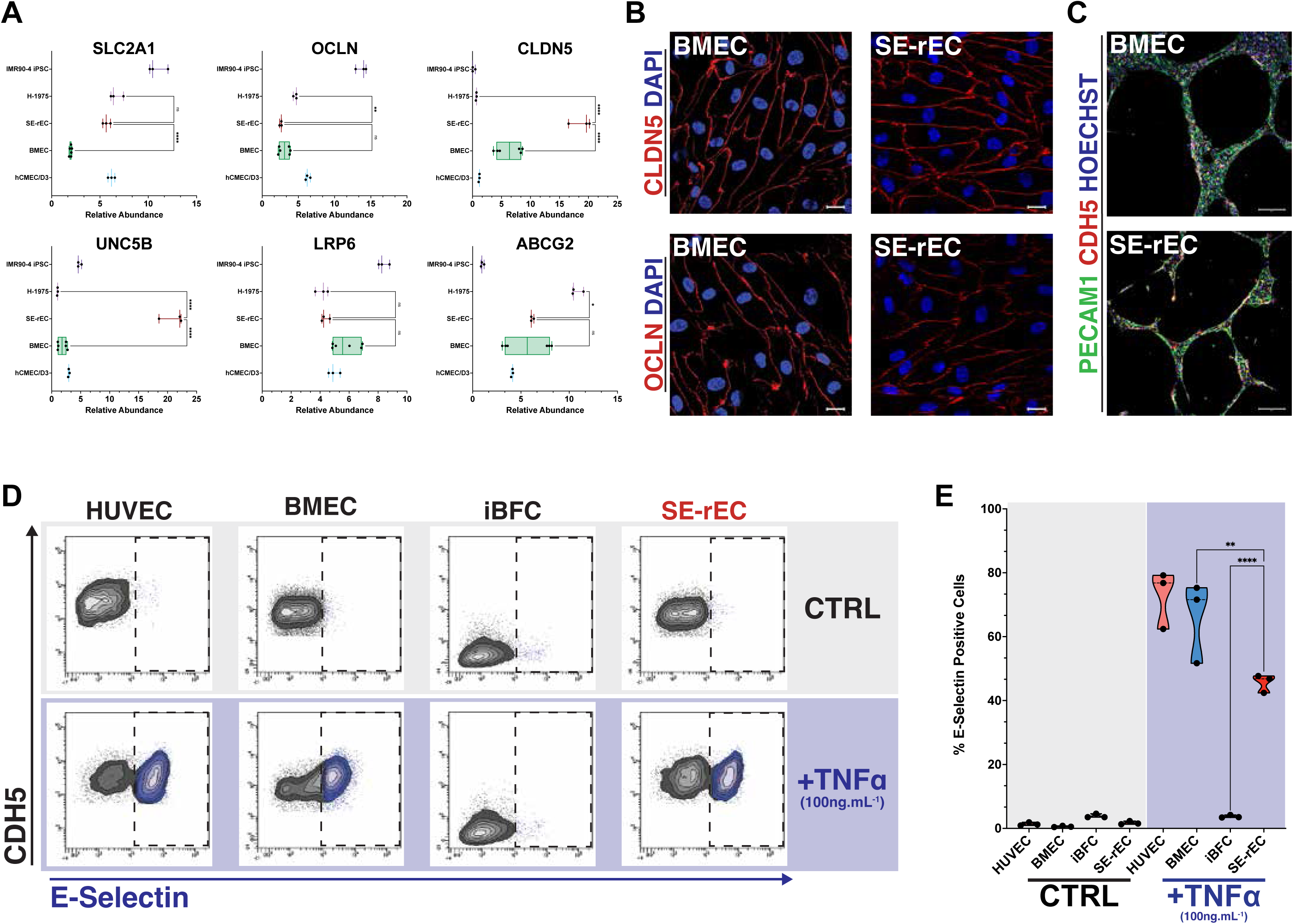
Reprogramming by SOX7 and ERG drives acquisition of functional vascular phenotype *in vitro*. **A.** Box and whisker plots demonstrating relative abundance of select brain-specific endothelial proteins in BMECs (N=6), hCMEC/D3 (N=3), SE-rECs endothelial samples (N=3), H-1975 epithelial samples (N=3), and IMR90-4 iPSC samples (N=3) **B.** Immunofluorescence for CLDN5(Red; top) and OCCLUDIN(Red; bottom) and DAPI (blue) in IMR90-4 derived SE-rECs and BMECs (scale bars=50µm; N=5 biological replicates). **C.** Tube formation assay showing vascular specific tube formation of both primary BMEC and SE-rECs on Matrigel with PECAM1(green), CDH5(red), DAPI(blue) (scale bars=100µm; N=5 biological replicates). **D.** Flow cytometry plots demonstrating E-Selectin and CDH5 protein expression in HUVEC, BMEC, Epi-iBFCs, and SE-rEC samples with and without stimulation by 100ng.ml-1 TNFα. **E.** Quantification of flow cytometry data from panel D showing percent of E-selectin positive cells across control and TNFα treated samples (N=3 biological replicates). *P-value<0.05; **<0.002; ***<0.0002; ****<0.0001.

The preceding analyses establish that SE-rECs possess a canonical vascular endothelial transcriptome and proteomic profile while also retaining a HOX transcript expression repertoire comparable to *in vivo* brain EC samples. This endothelial cellular identity, however, does not address whether SOX7, ERG reprogramming of neuroectoderm drives any regional, brain-specific endothelial program that defines the neurovascular unit *in vivo*. To interrogate this directly, we benchmarked the monoculture transcriptomes of the five *in vitro*populations (SE-rEC, primary BMEC, EEF-rEC, HUVEC, and iBFC reference) against the Siletti et al. adult-human-brain endothelial atlas^62^ using literature-curated, EC-specific regional marker gene sets^2,63,101,102^ (**Fig. 5A**). When quantified for forebrain enrichment (mean forebrain Z minus mean hindbrain Z), SE-rECs were found to be forebrain-biased (+0.68). The epithelial iBFC reference scored higher still (+0.86) and the inverse of EEF-rECs (−0.90) and HUVECs (−0.50), while primary BMEC demonstrated a near-neutral enrichment Z-score (−0.14) (**Fig. 5A**). Upon further investigation of this meta-analysis, SE-rECs displayed the strongest concordance with hippocampal and temporal cortex regions of the forebrain when compared to the other cell populations, whereas reprogrammed EEF-rECs and primary HUVECs were mapped toward the cerebellum and pons regions of the hindbrain (**Fig. 5B**). Notably, this regional forebrain bias in SE-rECs is retained despite the cells not clustering with *in vivo* brain-ECs in the bulk RNA-seq analysis. This indicates that SOX7 and ERG mediated reprogramming establishes an endothelial profile that preserves the forebrain regional identity from the neuroectodermal origin rather than defaulting to the hindbrain-skewed identity of ETS-factor reprogrammed or umbilical ECs.

**Figure 5:**
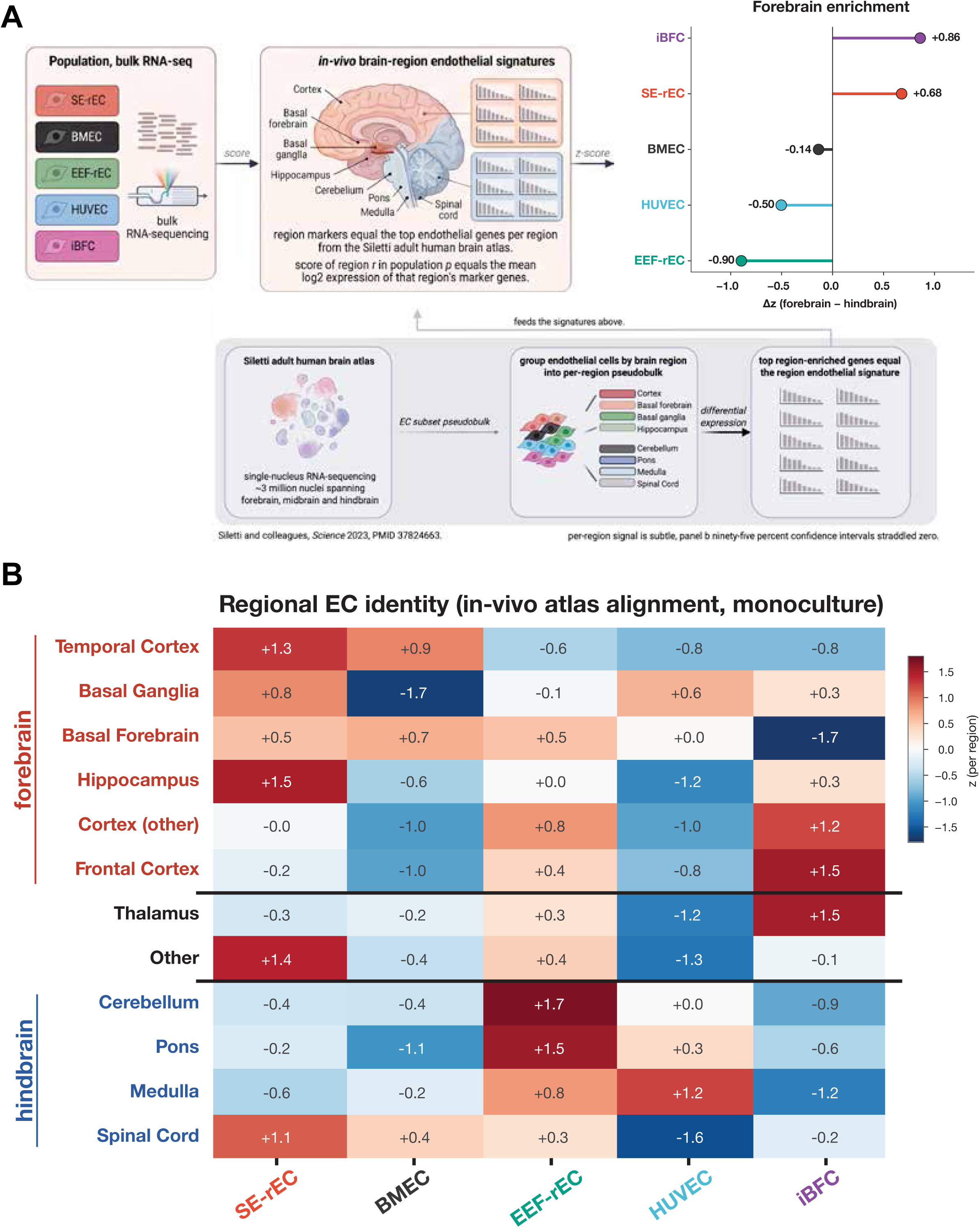
SOX7, ERG reprogramming yields a forebrain-biased endothelial transcriptome. **A.** Schematic of brain region-specific meta-analysis workflow. Per-region Z-score (across the five populations) of literature-curated EC regional-marker scores benchmarked against the Siletti adult-human-brain endothelial atlas. Forebrain enrichment index = mean(forebrain region Z) − mean(hindbrain region Z): iBFC +0.86, SE-rEC +0.68, BMEC −0.14, HUVEC −0.50, EEF-rEC −0.90. **B.** Heatmap highlighting regional bias across all cell populations. SE-rEC demonstrated highest correlation across the forebrain/cortical block (z-score: temporal cortex +1.3, hippocampus +1.5); EEF-rEC and HUVEC shift to the hindbrain. iBFC demonstrates high forebrain marker score.

To reinforce expression of a brain organotypic transcriptional profile in SE-rECs, we established a co-culture system by incorporating human primary astrocytes and brain pericytes that together are the major cell types which make up the BBB *in vivo*. The experiment was set up with ECs (HUVECs or SE-rECs) co-cultured directly with these astrocytes and pericytes to allow for direct contact between all cell types to recapitulate cell-cell interactions that occur in the BBB. After 5 days of co-culture, cells were phenotypically assessed by flow cytometry and microscopy which demonstrated the cellular composition of the cultures as well as the structural orientation of the cells. The results report that not only were all 3 cell types able to survive for 5 days in direct co-culture, but there was also a shift in cell composition ratio of the co-culture with the mean EC, astrocyte, pericyte ratio being closer to 2:1:2 respectively (**Fig. S5A**). Additionally, using phase microscopy, we report that by day 5 all 3 cell types were in direct cellular contact with one another, resulting in the formation of distinct structures protruding from an otherwise monolayer of cells (**Fig. S5B**). FACS isolation was then used to sequester SE-rECs from the co-cultures after which RNA was isolated for bulk RNA-seq and downstream bioinformatics analyses (**Fig. 6A**). The resulting principal component analysis demonstrated a clear clustering of co-culture EC samples away from the monocultured controls via PC2 whereas PC1 separated SE-rECs from the HUVEC controls (**Fig. 6B, S5C**). Initial transcriptomic analysis found that SE-rECs upregulate the expression of many vascular junction (e.g. *CLDN5*, *TJP1*) and brain enriched transporter genes (e.g. *SLC2A1*, *MFSD2A*) after five days in the *in vitro* co-culture. Conversely, this differential expression of organotypic transcripts was not observed in co-cultured HUVEC samples indicating limited microenvironmental attunement toward brain organotypic cell interactions (**Fig. 6C, D**). Genes known to affect vascular barrier permeability such as *ESAM*, *TJP1*, and *DLL4* were also among the subset of transcripts upregulated only in SE-rECs upon co-culture with these neural cell types (**Fig. 6C, D)**. To gain a better understanding of the underlying pathways that are modulated in SE-rECs after direct cellular contact with perivascular BBB cell types, we conducted a gene-set enrichment analysis incorporating all transcripts that were significantly upregulated (FDR-adjusted p-value<0.05) after 5 days in co-culture. Mapping these genes to functional cellular pathways revealed enrichment for processes such as “blood vessel development”, “tube morphogenesis” and “extracellular matrix organization” which demonstrated that these cells are undergoing membrane and structural changes which are in line with establishment of a BBB phenotype (**Fig. 6E**). Given its importance in the development and maintenance of the BBB, we sought to validate the transcript expression of *MFSD2A* as an indicator of brain organotypicity. MFSD2A protein was detected in co-cultured SE-rECs using confocal microscopy (**Fig. 7A**); with the highest expression detected in cells with close proximity to astrocytes and pericytes (**Fig. 7B**).

**Figure 6:**
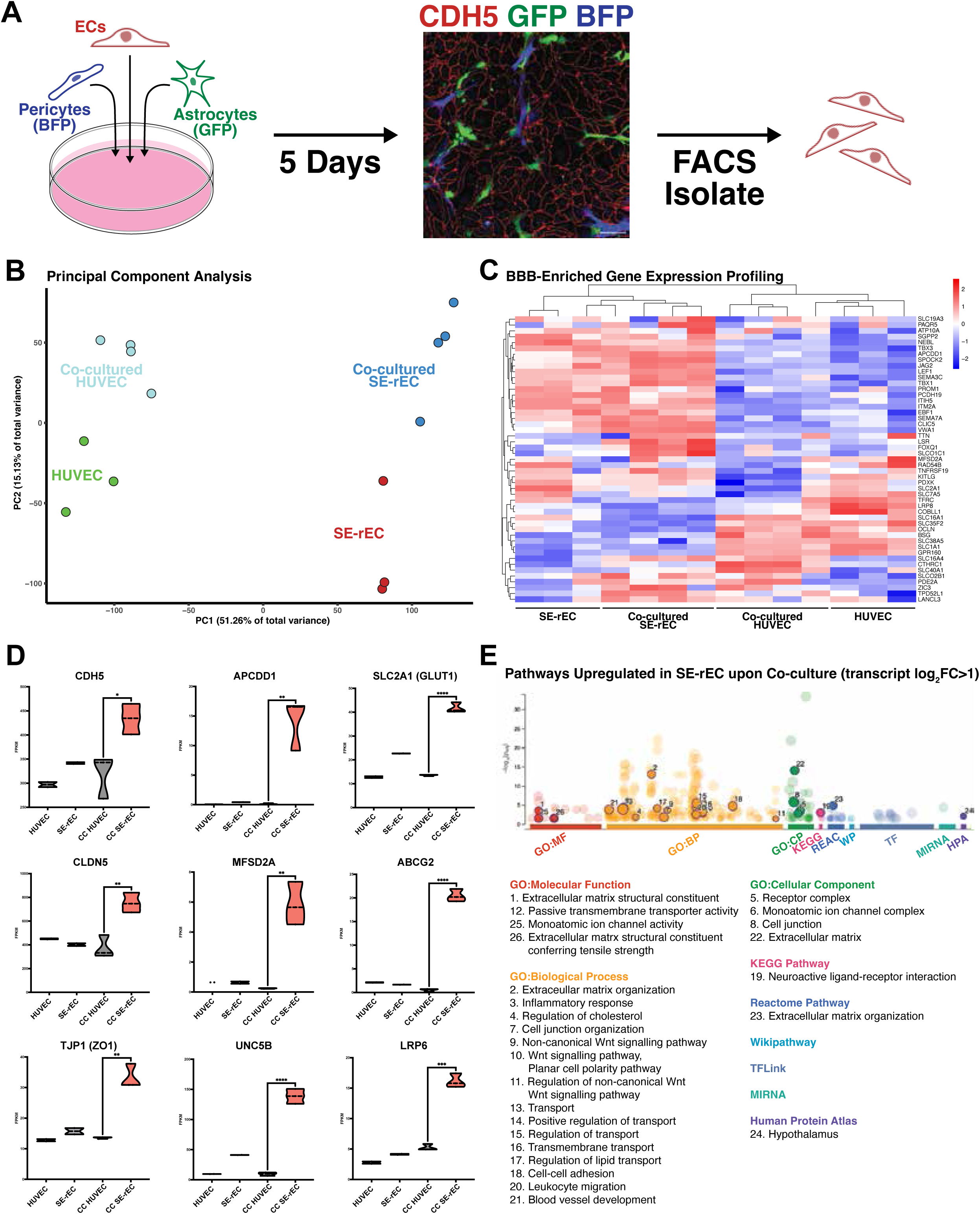
Direct co-culture of SE-rECs with Astrocytes and Pericytes in 2D drives acquisition of brain organotypicity. **A.** Schematic of the direct co-culture with example immunofluorescence of Astrocytes(GFP; Green), human brain Pericytes(BFP; Blue), and SE-rECs with CDH5(Red). (scalebar=100µm) **B.** Principal component analysis of HUVEC and SE-rEC global transcriptomes with and without 5 days of direct co-culture with human brain pericytes and astrocytes **C.** Heatmap denoting expression of select BBB-enriched endothelial transcripts previously described in Munji et al., 2020^108^ in all mono- and co-cultured HUVEC and SE-rEC samples **D.** Violin plots showing differential transcript expression of select BBB-specific genes in mono- and co-cultured HUVEC and SE-rEC samples. *P-value<0.05; **<0.002; ***<0.0002; ****<0.0001 **E.** Geneset enrichment analysis utilizing all genes upregulated in SE-rECs upon 5 days of direct co-culture (log_2_FC>1) demonstrating upregulation of specific gene ontology, KEGG, wikipathway, and human protein atlas pathways

**Figure 7:**
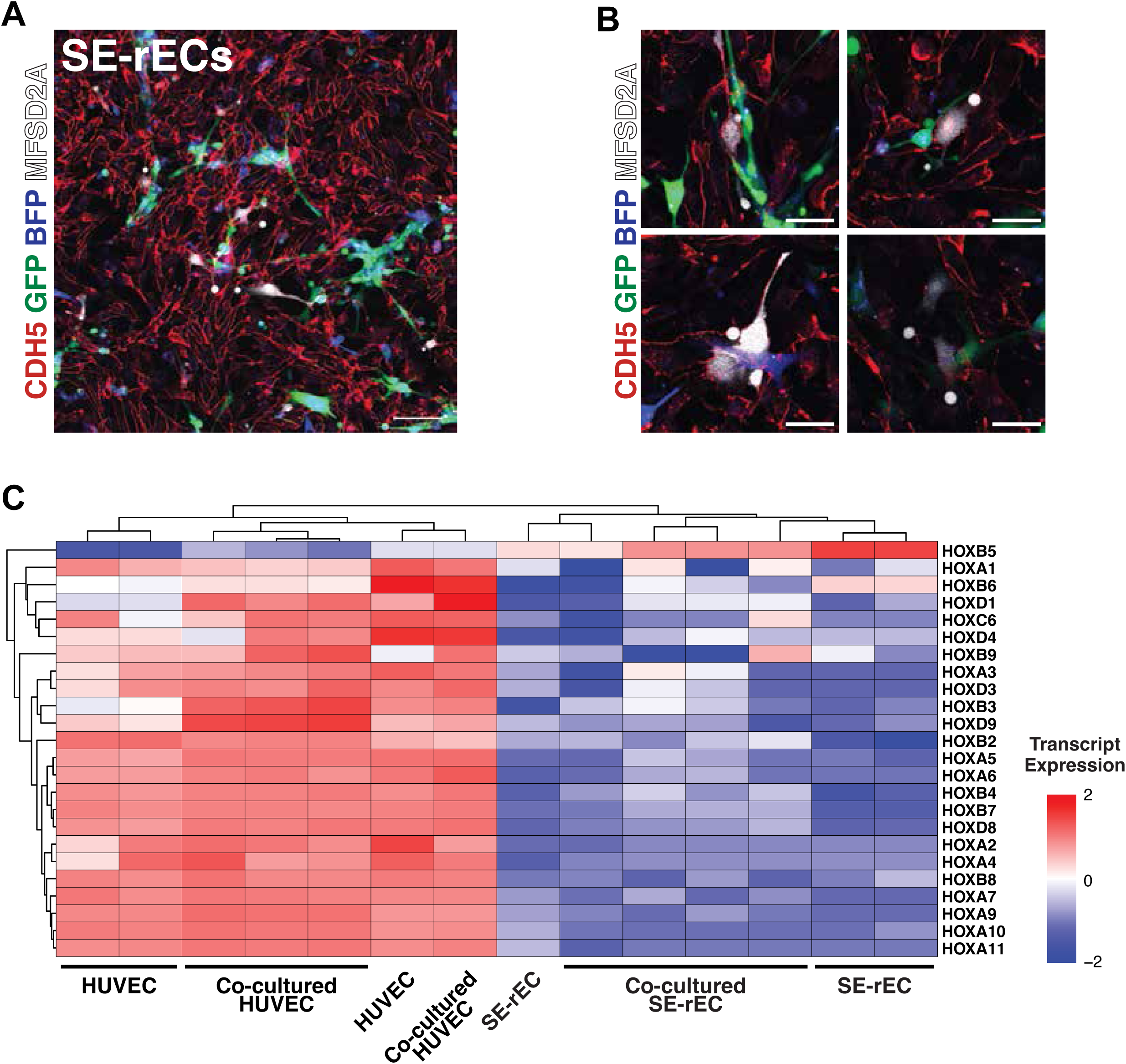
Direct co-culture promotes brain specific spatial honing of SE-rECs. **A.** Confocal microscopy images highlighting expression of MFSD2A (white) in SE-rECs labeled with CDH5 (red) after 5 days of co-culture with astrocytes (GFP; green) and pericytes (BFP; blue) (n=5 biological replicates; scale bars=100µm) **B.** Confocal microscopy images highlighting expression of MFSD2A (white) in SE-rECs labeled with CDH5 (red) after 5 days of co-culture with astrocytes (GFP; green) and pericytes (BFP; blue) (n= 5 biological replicates; scale bars=25µm) **C.** Heatmap of bulk RNAseq transcriptomes demonstrating low HOX gene expression in all SE-rEC samples even after co-culture with astrocytes and pericytes.

Following these initial transcriptomic analyses, we investigated the HOX gene expression profiles of HUVECs and SE-rECs with and without co-culture with astrocytes and pericytes. Expression of most HOX genes was low or absent in SE-rECs compared to HUVECs across both culture conditions(Fig. 7C). This suggests that our reprogrammed ECs have both retained HOX patterning from neuroectodermal embryonic origins and possess spatial recognition of the current microenvironment. Our results also demonstrate a complete lack of HOX specific adaptation to the neural vascular microenvironment by the lateral plate mesoderm-derived HUVECs.

Two-dimensional *in vitro* culture conditions do not adequately model many aspects of *in vivo* physiology as cells are plated on hard and flat surfaces, often limiting the amount of direct cellular interactions that can occur between neighboring cells. These conditions also usually lack flow-induced shear stress which is an essential aspect of modelling a vascular system. Perfusable organ-on-chip models have been widely utilized to circumvent these limitations and serve as a more functionally relevant *in vitro* BBB model. We chose to utilize the Mimetas OrganoPlate platform which allowed for both co-culture of EC and neural cell types in separate compartments and introduction of laminar flow to the ECs. To establish our organ-on-chip we first cultured either SE-rECs or primary BMECs on the inner periphery of a cylindrical chamber. This methodology results in a tube-like structure of ECs through which media can be flowed. Astrocytes and pericytes were then seeded in suspension in a parallel chamber that possesses a semi-permeable membrane. These neural cells would then migrate towards the EC chamber through a middle chamber of collagen gel separating the EC and neural cell compartments. This seeding strategy results in an endothelial vessel-like structure to which shear-stress can be applied that is also in contact with astrocytes and pericytes through a membrane allowing for direct cellular crosstalk (**Fig. 8A, B**).

**Figure 8:**
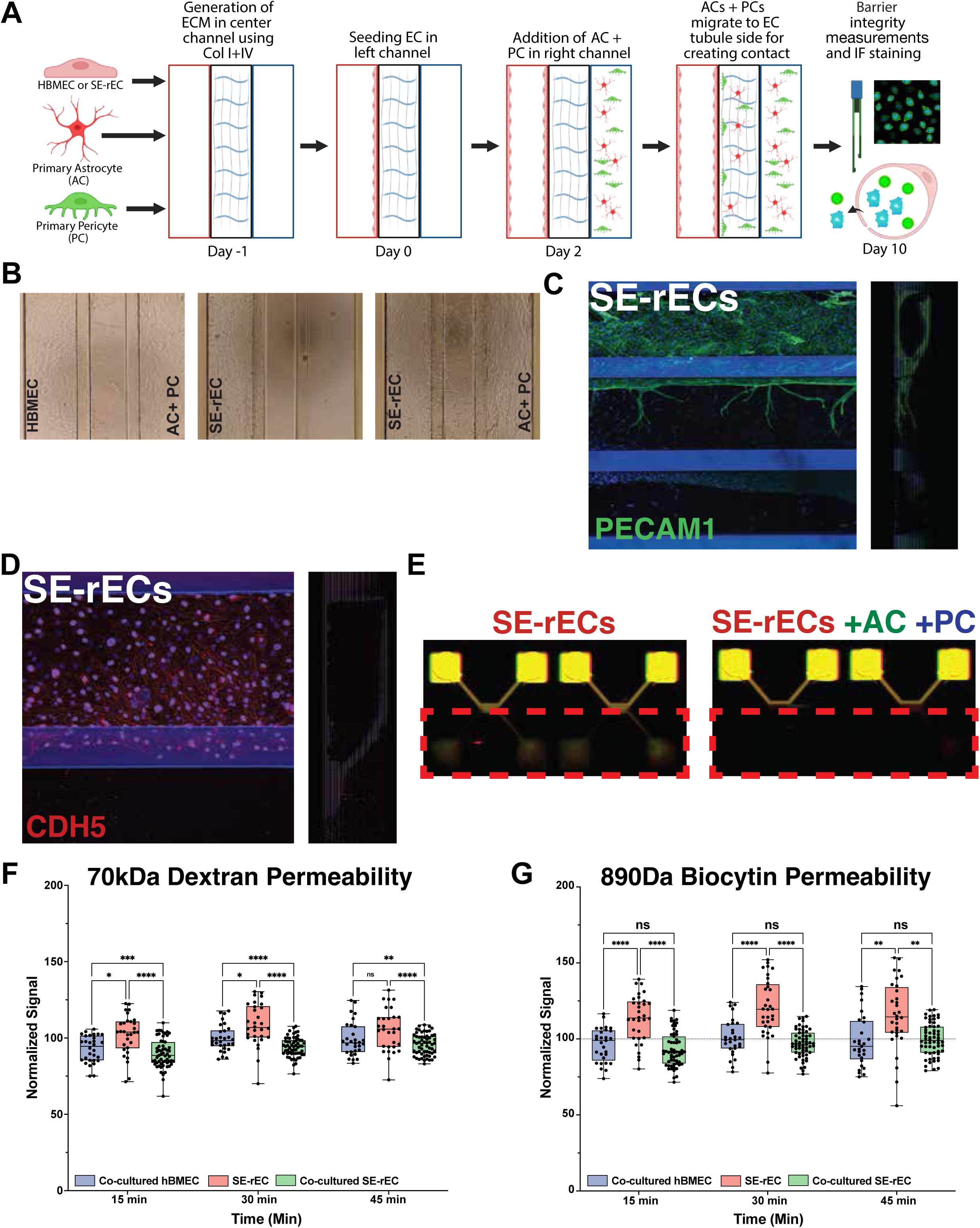
SE-rECs functionally adapt to 3-dimensional co-culture in an OrganoPlate device. **A.** Schematic of SE-rECs co-cultured with Astrocytes and Pericytes in the Mimetas OrganoPlate device **B.** Phase microscopy of Mimetas OrganoPlates seeded with human BMEC or SE-rECs, astrocytes, and pericytes. **C.** Immunofluorescence of SE-rECs co-cultured in Mimetas OrganoPlate device showing PECAM1(Green) and DAPI(Blue) **D.** Immunofluorescence of SE-rECs co-cultured in Mimetas OrganoPlate device showing CDH5(Red) and DAPI(Blue) **E.** SE-rEC monoculture (left) and co-cultured (right) in Mimetas OrganoPlate device demonstrating leakage of 70kD dextran tracer across middle ECM chamber **F.** Quantification of 70kD dextran signal in the bottom chamber of the Mimetas OrganoPlate device seeded with SE-rECs, Astrocytes, and Pericytes **G.** Quantification of 890D Biocytin signal in the bottom chamber of the Mimetas OrganoPlate device seeded with SE-rECs, Astrocytes, and Pericytes

After co-culture, expression of vascular barrier associated markers such as PECAM1 and CDH5 were demonstrated in the SE-rECs by immunofluorescence (IF) microscopy of the NVU-on-chip model system (**Fig. 8C, D)**. Subsequently, a high molecular weight tracer (70kDa dextran) was used to assess functional changes in vascular barrier phenotype (**Fig. 8E, F**). Co-cultured SE-rECs demonstrated significantly decreased permeability to dextran compared to the monocultured condition indicating an increase in barrier function. The barrier phenotype was not observed in cultures of astrocytes and pericytes alone. A similar trend of reduction was observed for the small-molecule tracer biocytin (≈890 Da; **Fig. 8G**).

## Discussion

Recent advancements in the development of *in vitro* models have allowed for the study of various functional characteristics of the BBB. However, current models are predicated on the accepted consensus that *in vitro* models of the BBB only need to capture a select function of interest such as tight junction mediated barrier formation or immune cell interaction. This has led to the use of some models that utilize cells which do not possess a vascular endothelial identity. However, to study a biological system as intricate and cell-specific as the BBB, one needs to adopt a model which is not only functionally congruent with the *in vivo* system, but is also comprised of cells harboring the correct cellular identities. This is especially true if the goal of such studies is to elucidate developmental mechanisms or develop targeted therapeutics which can both be very cell type dependent. Use of nonvascular cells may lead to critical misunderstandings regarding molecular pathways in homeostatic or pathophysiological states as well as dangerous off target effects from therapies designed to target a cell-specific marker. Generic ECs also lack many of the key junctional proteins or transporters that play major roles in drug transport which could hinder the development of efficacious therapies designed to navigate through the BBB.

Here, our work demonstrates the generation of bona-fide ECs from hPSC sources using *SOX7* and *ERG* constitutive expression which are transcriptomically, proteomically and functionally congruent to generic vascular ECs. These SE-rECs were able to demonstrate both angiogenic potential by forming tubes in Matrigel plugs and an EC-specific response to stimulation with inflammatory cytokines by upregulated E-selectin to the membrane in the presence of TNF-α. To further consolidate the cellular identity of SE-rECs, we investigated the expression of HOX genes across our dataset as brain ECs have been reported to express a very low level of all HOX family transcripts^11,12^. In cranial tissues, including the forebrain and its vasculature, HOX expression is largely absent, reflecting the developmental origin from HOX-negative cranial neural crest and anterior mesoderm^103,104^. Our meta-analysis supports previous reports as *in vivo* BMECs from human biopsies possessed negligible expression of almost all HOX transcripts, consistent with this developmental model. Prior *in vivo* and *in vitro* studies have suggested that HOX-encoded positional identity constrains endothelial heterogeneity and organotypicity^105,106^. Remarkably, SE-rECs exhibited a similar HOX-low transcript profile, clustering closely with i*n vivo* BMECs and distinct from *in vitro* BMECs that upregulate HOX gene expression in culture. This suggests that the reprogramming process did not induce ectopic HOX activation and that the neuroectodermal developmental programming of iBFCs may have preserved anterior cellular identity. The maintenance of this positional signature could be responsible for the partial BBB-like endothelial characteristics observed in SE-rECs.

A distinguishing feature of this reprogramming strategy emerged when SE-rECs were benchmarked against a human adult-brain endothelial atlas^62^. Unlike the permissive ETS-only reprogramming strategies, which yields a regionally undefined endothelium, SOX7/ERG conferred a forebrain-biased regional transcriptional program that was stronger than that of EEF-rECs or umbilical-vein endothelium. We interpret this as evidence that the SOXF factor SOX7, whose expression has been shown to be functionally required by brain ECs during development^85–89,107^, biases the reprogrammed cells toward the forebrain coordinates of their neuroectodermal origin rather than defaulting to the hindbrain-skewed or generic identity acquired through ETV2 induction. Importantly, this claim has specific boundaries. For instance, the forebrain program is a relative, regional bias rather than an absolute attainment of in-vivo brain-endothelial regional identity. The forebrain program installed by SOX7, ERG reprogramming is therefore best described as a relative, reprogramming bias caused by the transcription factors used (SOX, ETS vs ETS alone) rather than an absolute attainment of brain-endothelial regional identity. The conceptually important claim is therefore the conjunction of two properties the field has tended to treat separately: regional specificity (i.e. forebrain) and an authentic endothelial lineage identity.

Subsequently, we illustrated the use of organotypic microenvironmental cues to drive brain organotypicity in these hPSC-derived SE-rECs. Utilizing an *in vitro* co-culture system, SE-rECs were seeded in direct contact with both astrocytes and pericytes allowing for cellular crosstalk and tight junction formation that models the *in vivo* BBB. As a result of this molecular coaxing, SE-rECs which harbor a canonical vascular endothelial identity (CDH5^+^PECAM1^+^KDR^+^) upregulate expression of BBB specific transporters and junctional markers (e.g. *SLC2A1*, *MFSD2A*, *CLDN5*). Notably, upregulation of these BBB specific genes occurred only in SE-rECs, whereas HUVECs in the same co-culture system failed to demonstrate the same transcriptomic changes. Gene ontology pathways correlated with vasculature development and extracellular matrix organization were amongst the biological processes mapped to the upregulated genes uniquely in co-cultured SE-rECs; thereby representing transcriptomic evidence for the establishment of a vascular barrier phenotype. Under these 2D co-culture conditions, SE-rECs demonstrate retention of spatially restricted developmental HOX transcript expression patterning previously shown to be consistent with neuroectodermal origins. The fact that these changes were not observed in HUVECs under identical co-culture conditions underscores the importance of intrinsic deterministic HOX patterning aligned with a cell’s embryonic origin. This highly conserved and specific HOX patterning could play a major role in interpreting organ-specific signals but future studies will be needed to further investigate the mechanisms behind these results. SE-rECs, having been reprogrammed through this transcription-factor-mediated strategy that follows developmental spatiotemporal hierarchy, suggesting a transcriptional landscape conducive to the interpretation of astrocyte and pericyte-derived cell signaling. The co-culture-induced expression of MFSD2A, a lipid transporter critical to the formation and function of the BBB, demonstrates that these developmental pathways can be reinitiated *in vitro* through appropriate cell–cell interactions. Moreover, confocal microscopy revealing MFSD2A localization in SE-rECs in close proximity to astrocytes and pericytes provides spatial validation of paracrine and juxtacrine signaling specificity. This finding builds on the established knowledge given that MFSD2A expression *in vivo* is tightly dependent on perivascular astrocyte-derived WNT7A/B signaling^27^, establishing a possible mechanistic bridge between our direct co-culture system and developmental processes governing BBB specification during embryogenesis^101^.

Transitioning from 2D culture to the Mimetas OrganoPlate® platform provided an opportunity to interrogate the contribution of flow dynamics and 3D architecture to endothelial barrier function. Within this microfluidic device, SE-rECs were cultured on the periphery of a tube which led to the formation of a vascular microtube that could be continuously perfused^27^. This allowed the application of low-magnitude, bidirectional rocker-driven flow (∼2-5dyn/cm^2^). Functionally enhanced barrier integrity was demonstrated in SE-rECs co-cultured with astrocytes and pericytes in this system through a reduction in 70-kDa dextran permeability (p<0.05 between mean fluorescence intensities). SE-rECs were also found to decrease permeability to 890-Da biocytin when co-cultured. This demonstrates that BBB co-culture induced structural and functional maturation and this increased barrier phenotype was not observed in monoculture. The sensitivity of SE-rECs to the microenvironment could also reflect an intermediate state of differentiation that confers both adaptability and responsiveness, traits reminiscent of developing endothelium *in vivo*.

The establishment of a physiologically and transcriptomically relevant model of the NVU is a critical milestone in the study of the BBB and development of targeted therapeutics. This work represents a novel approach for the generation of hPSC-derived cells which harbor a vascular EC cellular identity which can be coaxed towards a brain specific phenotype and used in a 3-D organ-on-chip device. The ability for SE-rECs to upregulate expression of BBB specific surface markers, transporters, and WNT pathway-related genes in the presence of an NVU microenvironment makes them suitable for the study of various brain organotypic phenotypes. The advent of *in vitro* models utilizing hPSC lines which harbor disease specific mutations has aided in the study of cell specific roles in disease etiology as well as cell specific responses to therapies. In this context, SE-rECs derived from these disease specific hPSC lines could be used to study the vascular compartment of the BBB during different pathophysiological states. Our vascular organ-on-chip model could also ideally be used to screen targeted cellular therapies and small molecule drugs which need to traverse the BBB. Thus, we believe the multicellular *in vitro* model described in this work could eventually be used to advance our understanding of neurological diseases as well as in translational applications.

## Author contributions

T.M.L, U.A., D.A. and R.L. designed and conducted various aspects of the experiments and wrote and edited the manuscript. S.H. and T.M.L. analyzed sequencing data and performed computational analysis. A.J.R., A.C., J.W., F.G., V.A.M., S.S., and J.X. provided technical assistance. Z.R., S.R., I.M.P., J.G., R.W., F.C.B., and P.G. helped to formulate various aspects of the hypothesis and interpret the results. D.A. & R.L. directed and supervised all aspects of the study and manuscript. All authors edited the final manuscript.

## Acknowledgements

We thank Raul Chavez and the staff in the Genetic Resources Core Facility in Weill Cornell Medicine for technical assistance in various capacities. We also thank Chenghua Gu from Harvard Medical School for providing key reagents for this work. This work was supported by the National Institute of Health (#R61HL159949, #R33HL159949). T.M.L. and R.L. are also supported by the Ansary Stem Cell Institute, Starr Foundation TRI-Institution Stem Cell Initiative grants (Stem Cell Derivation Core, #2016-013).

## Data availability

Newly generated bulk RNA-sequencing data will be deposited in the Gene Expression Omnibus. Mass-spectrometry proteomics data will be deposited to the ProteomeXchange Consortium via the PRIDE partner repository. Previously published datasets re-analyzed here are listed under their original GEO accessions in the Methods.

---

**Figure S1**

**A.** Heatmap demonstrating expression of ETS and SOX genes across all in vitro iBFC and endothelial cell populations using previously published and newly generated datasets

**B.** Expression of SOX transcription factors found to be expressed across all in vitro endothelial samples from the meta-analysis in **Fig. S1**

**C.** Expression of ETS transcription factors found to be expressed across all in vitro endothelial samples from the meta-analysis in **Fig. S1**

**D.** Scree plot indicating the percent of explained variances denoted in each principal component of the analysis from **Fig. 1F**

**E.** Heatmap demonstrating expression of highest differentially expressed transcripts across PC2 of the principal component analysis from **Fig. 1F**

**Figure S2**

**A.** Pearson correlation analysis of the complete global transcriptomes from all samples from **Fig. 1F** demonstrating high correlation between all endothelial samples including SE-rECs according to total gene expression profile

**Figure S3**

**A.** Heatmap demonstrating expression of select canonical endothelial and epithelial marker genes across all samples

**B.** Violin plots showing relative abundance of select proteins across endothelial, epithelial, and hPSC samples

**Figure S4**

**A.** Heatmap demonstrating relative abundance of proteins driving molecular distance across PC2 in the proteomic principal component analysis

**B.** Heatmap demonstrating relative abundance of proteins associated with transcytosis across hPSC, epithelial and endothelial cell samples

**C.** Heatmap demonstrating relative abundance of all SLC proteins mapped across hPSC, epithelial and endothelial cell samples

**Figure S5**

**A.** Proportion of SE-rECs, Astrocytes, and Pericytes in culture after 5 days of co-culture at an original seeding density of 2.5:1:1

**B.** Phase microscopy images of HUVEC and SE-rECs in mono- and co-culture with Astrocytes and Pericytes. (scale bars=350µm)

**C.** Heatmap of the top 150 PC2 loading genes from **Fig. 6B** demonstrating expression in HUVEC and SE-rECs in mono- and co-culture with Astrocytes and Pericytes

## Methods

### Isolation of Human Umbilical Vein Endothelial Cells

Human umbilical vein endothelial cells (HUVECs) were de-identified and isolated in accordance with Weill Cornell Medicine IRB #0804009728R011 as described previously^109,110^. In short, Human umbilical cord samples were cannulated at each end flushed with HBSS to remove blood and identify leaks. Collagenase was then used to digest the tissue for 15 minutes at 37C. The cord was then flushed with HBSS to collect cells in a conical tube which was centrifuged at 1400rcf for 10 minutes. Cells were then resuspended in human EC medium (M199, 10% FBS, 50μg.ml^−1^ endothelial mitogen, and 100μg.ml^−1^ heparin) with penicillin and streptomycin to be plated onto 0.1% gelatin coated plates.

### Maintenance of Primary Human Cell Lines

HUVECs were initially maintained and expanded in human EC media following established protocol^109,110^. Brain microvascular endothelial cells were obtained from Sciencell Research Laboratories (cat. no. 1000) and were maintained following ScienCell guidelines. During the course of experiments, all endothelial cells were maintained in serum free endothelial media (DMEM/F12 Basal Media, 20% Knock-out Serum Replacement, 1ug.ml^−1^ insulin, 1x Glutamine Supplement, 1x MEM nonessential amino acids, 20mM HEPES buffer, 50ug.ml^−1^ heparin,10uM SB431542, 10ng.ml^−1^ recombinant human FGF-basic, 20ng.ml^−1^ recombinant human VEGF-A, 50ng.ml^−1^ recombinant human SCF) and seeded on 0.1% gelatin coated plates.

### Maintenance of hPSCs and differentiation to iBFCs

IMR90-4^111–114^ were obtained from WiCell.

These protocols were precisely followed as described previously^38,39,51^. All reagents were procured based on these protocols, and each step and duration for differentiation was followed rigorously. IMR90-4 hIPSCs were maintained between passages 20–42 on Matrigel (BD Biosciences, 354277) in mTeSR1 (STEMCELL Technologies). For differentiation, hPSCs were seeded onto Matrigel in mTeSR1 medium at 35,000cells/cm^2^ and allowed to expand for 3 days. Cultures were then switched to unconditioned medium (UM) consisting of DMEM/F12 Basal Media, 20% Knock-out Serum Replacement (Invitrogen), 1 mM L-glutamine (Sigma), 1x MEM nonessential amino acids (Invitrogen) for 6 days. EC medium consisting of human endothelial serum-free medium (hESFM; Gibco) supplemented with 20 ng.ml^−1^ FGF-basic (Peprotech), 1% platelet-poor plasma-derived bovine serum (PDS; Alfa Aesar), and 10 µM all-trans retinoic acid (Sigma, R2625) was then added for an additional 2 days. Cells were then dissociated with Accutase (Invitrogen) and plated onto 6-well polystyrene plates or 1.12cm^2^ Transwell-Clear permeable inserts (0.4 µm pore size) in EC media. Culture plates were coated with a mixture of Collagen IV (400 µg.ml−1; Sigma) and fibronectin (100 µg.ml−1; Sigma) in H_2_O for at least 30 min at 37°C, whereas inserts were incubated for a minimum of 4 h at 37°C. The resulting purified hPSC-derived iBFCs were then grown in EC medium for 24 h, after which RA was removed, and cells were cultured until the indicated experimental timepoints.

### Reprogramming iBFCs into SE-rECs

IMR90-4 or H6 iPSCs or H1 ESCs were differentiated according to the iBFC protocol as described above. Following the 6 days of differentiation in UM, cells were introduced to endothelial media for 2 days. On day 8 of the differentiation, the cells were dissociated using accutase (Corning), washed and transduced using lentiviral vectors for transcription factors SOX7 and ERG at an MOI of 10 and cultured on 0.1% gelatin in serum free endothelial medium at a 1:1 seeding density. After 7-14 days of expansion, cells were stained with PECAM1 (Biolegend), CDH5 (Biolegend), and EPCAM (Biolegend) after which PECAM1+CDH5+EPCAM-DAPI-cells were isolated using Fluorescence-activated cell sorting (FACS). These cells were cultured at a seeding density of 10,000-20,000 cell/cm^2^ on 0.1% gelatin with serum free endothelial medium being changed every 48 hours.

### Bulk RNA sequencing

Total RNA, greater than 100ng, from cultured cells was isolated in Qiagen Buffer RLT and purified using Qiagen RNeasy Mini Kit per manufacturer’s protocols. Agilent Technologies 2100 Bioanalyzer was used to assess the RNA quality. RNA Integrity Number (RIN) was used as a metric for RNA quality on a 1-to-10 scale (1 = fully degraded, 10 = intact). Samples with RIN of >7 were deemed acceptable quality for this study. RNA libraries were prepared and multiplexed using NEBNext® Ultra™ II Directional RNA Library Prep Kit for Illumina® (non-stranded and poly-A selection) and 10 nM of cDNA was used as the input for high-throughput sequencing via Illumina’s NovaSeq X Plus platform, producing 50 bp paired end reads. Previously published RNAseq data for iBFCs and various EC and epithelial cell types were downloaded from the Gene Expression Omnibus (GEO Accession numbers: GSE40291, GSE57662, GSE131039, GSE137786, GSE82207, GSE129290, GSE122588, GSE97100, GSE97324, GSE73721, GSE97575, GSE108012, GSE151976, GSE157852 and GSE126449) which were then processed in the same way as for the bulk RNAseq libraries prepared for the experiment.

Sample files were checked for sequence quality (FastQC v0.11.5) and processed using the Digital Expression Explorer 2 (DEE2)^115^ workflow. Adapter trimming was performed with Skewer (v0.2.2)^116^ using default parameters of a 3 base-pair minimum overlap for adapter detection and a maximum error ratio of 0.1. Further quality control done with Minion, part of the Kraken package^117^ filtering out read with low average Phred scores (<20). The resultant filtered reads were mapped to human reference genome GRCh38 using STAR aligner^118^ and gene-wise expression counts generated using the “-quantMode GeneCounts” with all parameters at their default values. Mapping quality parameters included a maximum number of mismatches allowed set to 10, a mismatch-to-read-length ratio of 1.0, and only uniquely mapped reads (MAPQ = 255) were retained for downstream analyses. After further filtering and quality control, R package edgeR^119^ was used to calculate TMM normalization factors. Batch effects between libraries was removed using the “removeBatchEffect” function in Limma (v3.65.3)^120^. Fragments Per Kilobase of transcript per Million mapped reads (FPKM) and Log2 counts per million (logCPM) matrices were quantified using these factors to normalize for library size. R functions prcomp and get_eigenvalue were used on normalized Log2 CPM values to compute principal components. R function cor was used to compute Pearson correlation. The ggplot2 R package was used to visualize all PCA plots and correlation plots. The pheatmap package was used to create dendrograms and visualize heatmaps and correlation plots. Dendrograms associated with heatmaps were created using pheatmap’s cluster_method parameter set to average Euclidean distance between samples or transcripts. Gene enrichment analysis was run using WebGestalt^121^. The complete bulk RNAseq meta-analysis contained 195 samples across 27 libraries. These included *in vivo* samples (N=11 brain endothelial samples isolated from patients) as well as *in vitro* samples (N=53 endothelial samples from various tissues, N=3 choroid plexus samples, N=107 iBFC samples, N=5 bronchial epithelial samples, N=8 iPSC-derived colon organoid samples, and N=8 SE-rEC samples).

### Endothelial inflammatory response assay

HUVEC, BMEC, as well as IMR90-4 derived iBFCs and rECs were cultured in serum free media until they reached confluent monolayer on plastic tissue culture plates coated with 0.1% gelatin (ECs) or a mixture of Collagen IV (400 µg.ml−1) and fibronectin (100 µg.ml−1) (iBFCs). 100ng ml^-1^ of Tumor Necrosis Factor alpha (TNFα) was added to culture media and allowed to stimulate cells for 3 hours. Cells were then washed, fixed, and assayed for E-Selectin (CD62E) by confocal microscopy and flow cytometry. One-way ANOVA using a normal (gaussian) distribution assumption was employed to compare the mean percentages of E-selection positively marked cells across samples in the control or TNFα treated conditions. N=3 biological replicates per cell type per condition; *P-value<0.05; **<0.002; ***<0.0002; ****<0.0001.

### Immunofluorescence

#### Paraformaldehyde Fixed Samples

Cells were fixed with 4% paraformaldehyde for 15 minutes prior to staining. To prevent non-specific binding of the primary antibody, samples were blocked with 5% horse serum in PBS solution for 60 minutes. Cells were then washed 3 times in PBS and dilute primary antibody in 5% horse serum was added according to manufacturer’s recommended concentration and left overnight at 4°C. If the protein was intracellular, permeabilization using 0.2% Triton X-100 was necessary during block and staining with primary antibody. After staining with primary antibody, cells were incubated with secondary antibody in 5% horse serum for 1 hour at room temperature on a shaker. Following a thorough wash to remove unbound secondary antibody, DAPI was used at 1 µg.ml^−1^ to stain nuclei.

#### Methanol Fixed Samples

Cells were fixed with 95% methanol for 30 minutes at -20°C and then quenched in Acetone for 5 minutes at room temperature. Cells were then washed 3 times in PBS and dilute primary antibody in 5% horse serum was added according to manufacturer’s recommended concentration and left overnight at 4°C. After staining with primary antibody, cells were incubated with secondary antibody in 5% horse serum for 1 hour at room temperature on a shaker. Following a thorough wash to remove unbound secondary antibody, DAPI was used at 1 µg.ml^−1^ to stain nuclei.

#### Live Samples

Human IgG antibody was used at 1:50 in PBS with 0.5% BSA and 2 mM EDTA to block Fc receptors prior to staining. The blocked cells were stained for 30 minutes at 4°C with fluorochrome-conjugated antibodies according to the manufacturer’s recommendations. Stained cells were washed in PBS to prevent saturation of the fluorescent dye during imaging.

All immunofluorescence imaging was performed using a Zeiss 710 META confocal microscope.

### Flow cytometry

Before staining, F_c_ receptors were blocked with human IgG antibody at 1:50 (Biolegend) in PBS (pH 7.2) containing 0.5% BSA (Fraction V) and 2mM EDTA for 10min at 4 °C. For cell surface staining, samples were stained for 30min at 4°C with fluorochrome-conjugated antibodies according to the manufacturer’s recommendation. Stained cells were washed in blocking buffer and fixed in 1% PFA in PBS (pH 7.2) with 2mM EDTA for flow analysis or resuspended in PBS (pH 7.2) with 2mM EDTA and 1 μg.ml^−1^ DAPI (Biolegend) for sorting. Samples were analysed using a FACS ARIA II SORP and/or FACS ARIA FUSION SORP (BD Biosciences). Data were collected and analyzed using FACs DIVA v9.7 software (BD Biosciences). All gating was determined using unstained controls and fluorescence minus one strategy. A minimum of N=3 biological replicates were used for all quantitative flow cytometry experiments.

### Tubulogenesis assay

IMR90-4 derived SE-rECs, and primary BMECs were seeded at 40,000cells/cm^2^ on fully polymerized Matrigel (Corning) for 24hours after which live staining was conducted using anti-human PECAM1, CDH5, and Hoechst following the live samples protocol detailed above. Imaging was performed using a Zeiss 710 META confocal microscope. N=5 biological replicates per cell type were used for this experiment.

### TMT Labeling Quantitative Proteomic Analysis

Samples for protein analysis were prepared essentially as previously described^122,123^. Proteomes were extracted using a buffer containing 200 mM EPPS pH 8.5, 8M urea, and protease inhibitors. Following lysis, each sample was reduced with 5 mM TCEP. Cysteine residues were alkylated using 10 mM iodoacetimide for 20 minutes at RT in the dark. Excess iodoacetimide was quenched with 10 mM DTT. 100 µg of each proteome was precipitated and re-solubilized in 200 mM EPPS pH 8.5. Samples were digested with Lys-C (1:50) overnight at RT and subsequently with trypsin (1:100) for 6 hours at 37°C. Anhydrous acetonitrile was added to each sample to achieve a final concentration of 33% acetonitrile. 50 µg of peptides from each sample were labeled with TMTPro reagents (ThermoFisher Scientific) for 2 hours at room temperature. Labeling reactions were quenched with 0.5% hydroxylamine and acidified with formic acid. Acidified peptides were combined and desalted by Sep-Pak (Waters). Proteome data were collected on an Orbitrap Eclipse mass spectrometer (ThermoFisher Scientific) with FAIMS enabled coupled to a Proxeon EASY-nLC 1200 LC pump (ThermoFisher Scientific). A minimum of N=3 biological replicates were used per sample type.

### Two-dimensional neurovascular-unit co-culture

For direct co-culture, FACS-purified SE-rECs or HUVECs were seeded together with primary human astrocytes and primary human brain pericytes (Sciencell Technologies) at a [2.5:1:1] endothelial:astrocyte:pericyte ratio for 5 days to permit direct cell–cell contact. The endothelial compartment was then re-isolated by fluorescence-activated cell sorting (EPCAM−PECAM1+CDH5+) and processed for bulk RNA sequencing as described below. Co-culture composition and purity were confirmed by flow cytometry (**Fig. S5**).

### Three-dimensional neurovascular-unit-on-chip and permeability assays

A 3D microfluidic blood–brain barrier (BBB) model was established using OrganoPlate 3-lane-64 devices (MIMETAS). Each chip contained an extracellular matrix (ECM) gel in the central channel, an endothelial tubule in the right channel, and a mixture of primary astrocytes and pericytes seeded in the left channel, which migrated toward the endothelial compartment during culture on a bidirectional OrganoFlow rocker.

On day −1, the middle channel was filled with a collagen I/collagen IV mixture (4:1; each at 5 mg/mL), incubated at 37 °C for 15 minutes, and then overlaid with PBS to prevent desiccation. The right channel inlet was coated overnight with fibronectin (10 µg/mL) and collagen IV (10 µg/mL). On day 0, endothelial cells were dissociated and resuspended at 20,000 cells/µL. A total of 40,000 cells were introduced into the right channel outlet of each chip. Plates were placed at a 75° angle (manufacturer stand) and incubated at 37 °C for 6 hours to allow cells to attach along the ECM boundary. Outlets were then replenished with endothelial medium, and chips were transferred to the OrganoFlow rocker alternating between −7° and +7° every 8 minutes. Within 24–48 hours, endothelial cells formed an open tubular structure. On day 1, the left inlet was coated with Matrigel. Primary astrocytes were resuspended in AM-PM medium (4:1 AM:PM) at 7,500 cells/µL, and primary pericytes were prepared similarly at 5,000 cells/µL. Cells were mixed at a 1:1 astrocyte:pericyte ratio, yielding 7,500 astrocytes and 5,000 pericytes per chip, and seeded through the left channel outlet. Plates rested for 4 hours at 37 °C before outlet wells were replenished with AM-PM medium and returned to rocker-based culture until day 10. Media was replaced every other day. Beginning on day 6, endothelial channels received standard EC medium lacking bFGF and VEGF.

On day 10, 70-kDa dextran (0.5 µg/mL) and biocytin (0.5 µg/mL) were added to the right inlet and outlet wells (40 µL and 30 µL, respectively). Wells of the ECM inlet, left inlet, and left outlet were filled with 20 µL tracer-free medium. Plates were placed on the rocker and imaged every 15 minutes using the LICOR system. Fluorescence intensities were quantified on the Odyssey SA. Permeability was expressed as normalized tracer intensity in the central ECM well. ECM-only chips served as controls for passive leakage. Statistical analysis used two-way ANOVA.

### Brain-region and organ endothelial-identity scoring

Regional endothelial identity was quantified using literature-curated, endothelial-cell-specific regional marker sets benchmarked against the Siletti adult-human-brain single-nucleus endothelial atlas; region- and organ-EC markers were curated from Ben-Zvi et al., Daneman et al., Vanlandewijck et al., Yang et al. and Siletti et al. For each population a per-region identity score was computed as the mean normalised log2 expression of that region’s curated markers and converted to a per-region Z-score across populations; the forebrain enrichment index was defined as the mean forebrain-region Z minus the mean hindbrain-region Z. Replicate-level statistical validation of this index is described below. Forebrain identity index and replicate-level statistics Bulk FPKM matrices were log2(FPKM + 1)-transformed and z-scored per gene across samples. A per-replicate forebrain identity index was defined as the mean z of a curated forebrain/cortical endothelial marker set (EFNB2, DLL4, HEY1, HEY2, GJA5, NOTCH4, NRP1, BMX, UNC5B, LSR, ITM2A, SPOCK2, APCDD1, SLC7A5, LEF1, AXIN2, TCF7L2, FZD4, NKD1) minus the mean z of a hindbrain endothelial marker set (MGP, IGFBP7, S100A16, BGN, NOTCH3, COL4A2, SPARCL1). Population means are reported with 95% confidence intervals from 5,000 bootstrap resamples over biological replicates. Between-population comparisons used one-sided Mann–Whitney U tests with Cliff’s δ effect sizes. Signature specificity was assessed against a permutation null of 20,000 random, size-matched marker-set pairs drawn from genes detected in ≥50% of samples. Reproducibility was tested in an independent SE-versus-EEF replicate cohort. Replicate counts: SE-rEC n = 8, in-vivo brain EC n = 11, primary BMEC n = 2, EEF-rEC n = 4, HUVEC n = 14, iBFC n = 65, ETV2-iEC n = 3. Analyses used numpy, pandas, scipy and matplotlib.

### Virus production

Human SOX7 and ERG lentiviral particles were generated and tittered as described in Ginsberg *et al.*^124^, with the following modifications. 293T Lenti-X cells (passage 8–10; subconfluent) were used to produce lentiviral particles using Lenti-X packaging single-shot system, following the manufacturer’s protocol. Supernatant was filtered through a 0.45 μm PES filter attached to a 30 ml syringe and concentrated using the Lenti-X concentrator according to the manufacturer’s protocol. Concentrate from a 100 mm plate of 293T cells was resuspended in 100 μl of lentivirus storage buffer and stored at −80 °C for up to 1 year.

## References

1 Reese, T. S. & Karnovsky, M. J. Fine structural localization of a blood-brain barrier to exogenous peroxidase. J Cell Biol 34, 207–217, doi:10.1083/jcb.34.1.207 (1967).

2 Vanlandewijck, M., He, L., Andaloussi Mäe, M., Andrae, J., Ando, K., Del Gaudio, F., Nahar, K., Lebouvier, T., Laviña, B., Gouveia, L., Sun, Y., Raschperger, E., Segerstolpe, Å., Liu, J., Gustafsson, S., Räsänen, M., Zarb, Y., Mochizuki, N., Keller, A., Lendahl, U. & Betsholtz, C. A molecular atlas of cell types and zonation in the brain vasculature. Nature 554, 475–480, doi:10.1038/nature25739 (2018).

3 Morita, K., Sasaki, H., Furuse, M. & Tsukita, S. Endothelial claudin: Claudin-5/TMVCF constitutes tight junction strands in endothelial cells. Journal of Cell Biology 147, 185–194, doi:10.1083/jcb.147.1.185 (1999).

4 Kniesel, U. & Wolburg, H. Tight junctions of the blood-brain barrier. Cell Mol Neurobiol 20, 57– 76, doi:10.1023/a:1006995910836 (2000).

5 Tsukita, S. & Furuse, M. The Structure and Function of Claudins, Cell Adhesion Molecules at Tight Junctions. Annals of the New York Academy of Sciences 915, 129–135, doi:10.1111/j.1749-6632.2000.tb05235.x (2006).

6 Luissint, A.-C., Artus, C., Glacial, F., Ganeshamoorthy, K. & Couraud, P.-O. Tight junctions at the blood brain barrier: physiological architecture and disease-associated dysregulation. Fluids and Barriers of the CNS 9, 23, doi:10.1186/2045-8118-9-23 (2012).

7 Greene, C., Hanley, N. & Campbell, M. Claudin-5: Gatekeeper of neurological function. Fluids and Barriers of the CNS 16, 1–15, doi:10.1186/s12987-019-0123-z (2019).

8 Engelhardt, B. Development of the blood-brain barrier. Cell and Tissue Research 314, 119–129, doi:10.1007/s00441-003-0751-z (2003).

9 Risau, W. & Wolburg, H. Development of the blood-brain barrier. Trends in Neurosciences 13, 174–178, 10.1016/0166-2236(90)90043-A (1990).

10 Bär, T. The vascular system of the cerebral cortex. Adv Anat Embryol Cell Biol 59, I–vi,1–62, doi:10.1007/978-3-642-67432-7 (1980).

11 Lippmann, E. S., Williams, C. E., Ruhl, D. A., Estevez-Silva, M. C., Chapman, E. R., Coon, J. J. & Ashton, R. S. Deterministic HOX patterning in human pluripotent stem cell-derived neuroectoderm. Stem Cell Reports 4, 632–644, doi:10.1016/j.stemcr.2015.02.018 (2015).

12 Molino, Y., Jabès, F., Bonnet, A., Gaudin, N., Bernard, A., Benech, P. & Khrestchatisky, M. Gene expression comparison reveals distinct basal expression of HOX members and differential TNF-induced response between brain- and spinal cord-derived microvascular endothelial cells. J Neuroinflammation 13, 290, doi:10.1186/s12974-016-0749-6 (2016).

13 Ma, X. J., Cheng, J. W., Zhang, J., Liu, A. J., Liu, W., Guo, W., Shen, F. M. & Lu, G. C. E-selectin deficiency attenuates brain ischemia in mice. CNS Neurosci Ther 18, 903–908, doi:10.1111/cns.12000 (2012).

14 Kisucka, J., Chauhan, A. K., Zhao, B.-Q., Patten, I. S., Yesilaltay, A., Krieger, M. & Wagner, D. D. Elevated levels of soluble P-selectin in mice alter blood-brain barrier function, exacerbate stroke, and promote atherosclerosis. Blood 113, 6015–6022, doi:10.1182/blood-2008-10-186650 (2009).

15 Ballabh, P., Braun, A. & Nedergaard, M. The blood-brain barrier: an overview: structure, regulation, and clinical implications. Neurobiol Dis 16, 1–13, doi:10.1016/j.nbd.2003.12.016 (2004).

16 Muoio, V., Persson, P. B. & Sendeski, M. M. The neurovascular unit – concept review. Acta Physiologica 210, 790–798, 10.1111/apha.12250 (2014).

17 Liebner, S., Dijkhuizen, R. M., Reiss, Y., Plate, K. H., Agalliu, D. & Constantin, G. Functional morphology of the blood-brain barrier in health and disease. Acta Neuropathol 135, 311–336, doi:10.1007/s00401-018-1815-1 (2018).

18 Profaci, C. P., Munji, R. N., Pulido, R. S. & Daneman, R. The blood-brain barrier in health and disease: Important unanswered questions. J Exp Med 217, doi:10.1084/jem.20190062 (2020).

19 Abbott, N. J., Rönnbäck, L. & Hansson, E. Astrocyte–endothelial interactions at the blood–brain barrier. Nature Reviews Neuroscience 7, 41–53, doi:10.1038/nrn1824 (2006).

20 Banks, W. A., Kovac, A. & Morofuji, Y. Neurovascular unit crosstalk: Pericytes and astrocytes modify cytokine secretion patterns of brain endothelial cells. Journal of Cerebral Blood Flow & Metabolism 38, 1104–1118, doi:10.1177/0271678X17740793 (2017).

21 Zlokovic, B. V. The blood-brain barrier in health and chronic neurodegenerative disorders. Neuron 57, 178–201, doi:10.1016/j.neuron.2008.01.003 (2008).

22 Daneman, R. The blood–brain barrier in health and disease. Annals of Neurology 72, 648–672, 10.1002/ana.23648 (2012).

23 Abbott, N. J., Revest, P. A. & Romero, I. A. Astrocyte-endothelial interaction: physiology and pathology. Neuropathology and Applied Neurobiology 18, 424–433, 10.1111/j.1365-2990.1992.tb00808.x (1992).

24 Bonkowski, D., Katyshev, V., Balabanov, R. D., Borisov, A. & Dore-Duffy, P. The CNS microvascular pericyte: pericyte-astrocyte crosstalk in the regulation of tissue survival. Fluids and Barriers of the CNS 8, 8, doi:10.1186/2045-8118-8-8 (2011).

25 Dohgu, S., Takata, F., Yamauchi, A., Nakagawa, S., Egawa, T., Naito, M., Tsuruo, T., Sawada, Y., Niwa, M. & Kataoka, Y. Brain pericytes contribute to the induction and up-regulation of blood– brain barrier functions through transforming growth factor-β production. Brain Research 1038, 208–215, 10.1016/j.brainres.2005.01.027 (2005).

26 Hori, S., Ohtsuki, S., Hosoya, K.-i., Nakashima, E. & Terasaki, T. A pericyte-derived angiopoietin-1 multimeric complex induces occludin gene expression in brain capillary endothelial cells through Tie-2 activation in vitro. Journal of Neurochemistry 89, 503–513, 10.1111/j.1471-4159.2004.02343.x (2004).

27 Daneman, R., Agalliu, D., Zhou, L., Kuhnert, F., Kuo, C. J. & Barres, B. A. Wnt/β-catenin signaling is required for CNS, but not non-CNS, angiogenesis. Proceedings of the National Academy of Sciences 106, 641–646, doi:10.1073/pnas.0805165106 (2009).

28 Sandoval, K. E. & Witt, K. A. Blood-brain barrier tight junction permeability and ischemic stroke. Neurobiology of Disease 32, 200–219, 10.1016/j.nbd.2008.08.005 (2008).

29 Stamatovic, S. M., Johnson, A. M., Keep, R. F. & Andjelkovic, A. V. Junctional proteins of the blood-brain barrier: New insights into function and dysfunction. Tissue Barriers 4, e1154641, doi:10.1080/21688370.2016.1154641 (2016).

30 Lochhead, J. J., Yang, J., Ronaldson, P. T. & Davis, T. P. Structure, Function, and Regulation of the Blood-Brain Barrier Tight Junction in Central Nervous System Disorders. Frontiers in Physiology 11, doi:10.3389/fphys.2020.00914 (2020).

31 Ayloo, S. & Gu, C. Transcytosis at the blood-brain barrier. Curr Opin Neurobiol 57, 32–38, doi:10.1016/j.conb.2018.12.014 (2019).

32 Pulgar, V. M. Transcytosis to Cross the Blood Brain Barrier, New Advancements and Challenges. Frontiers in Neuroscience 12, doi:10.3389/fnins.2018.01019 (2019).

33 Yang, A. C., Stevens, M. Y., Chen, M. B., Lee, D. P., Stähli, D., Gate, D., Contrepois, K., Chen, W., Iram, T., Zhang, L., Vest, R. T., Chaney, A., Lehallier, B., Olsson, N., du Bois, H., Hsieh, R., Cropper, H. C., Berdnik, D., Li, L., Wang, E. Y., Traber, G. M., Bertozzi, C. R., Luo, J., Snyder, M. P., Elias, J. E., Quake, S. R., James, M. L. & Wyss-Coray, T. Physiological blood–brain transport is impaired with age by a shift in transcytosis. Nature 583, 425–430, doi:10.1038/s41586-020-2453-z (2020).

34 Zhang, W., Liu, Q. Y., Haqqani, A. S., Leclerc, S., Liu, Z., Fauteux, F., Baumann, E., Delaney, C. E., Ly, D., Star, A. T., Brunette, E., Sodja, C., Hewitt, M., Sandhu, J. K. & Stanimirovic, D. B. Differential expression of receptors mediating receptor-mediated transcytosis (RMT) in brain microvessels, brain parenchyma and peripheral tissues of the mouse and the human. Fluids and Barriers of the CNS 17, 47, doi:10.1186/s12987-020-00209-0 (2020).

35 Daneman, R., Zhou, L., Agalliu, D., Cahoy, J. D., Kaushal, A. & Barres, B. A. The mouse blood-brain barrier transcriptome: a new resource for understanding the development and function of brain endothelial cells. PLoS One 5, e13741, doi:10.1371/journal.pone.0013741 (2010).

36 Nishihara, H., Gastfriend, B. D., Kasap, P., Palecek, S. P., Shusta, E. V. & Engelhardt, B. Differentiation of human pluripotent stem cells to brain microvascular endothelial cell-like cells suitable to study immune cell interactions. STAR Protoc 2, 100563, doi:10.1016/j.xpro.2021.100563 (2021).

37 Matsuo, K., Engelhardt, B. & Nishihara, H. Differentiation of Human Induced Pluripotent Stem Cells to Brain Microvascular Endothelial Cell-Like Cells with a Mature Immune Phenotype. J Vis Exp, doi:10.3791/65134 (2023).

38 Lippmann, E. S., Azarin, S. M., Kay, J. E., Nessler, R. A., Wilson, H. K., Al-Ahmad, A., Palecek, S. P. & Shusta, E. V. Derivation of blood-brain barrier endothelial cells from human pluripotent stem cells. Nature Biotechnology 30, 783, doi:10.1038/nbt.2247 https://www.nature.com/articles/nbt.2247#supplementary-information (2012).

39 Lippmann, E. S., Al-Ahmad, A., Azarin, S. M., Palecek, S. P. & Shusta, E. V. A retinoic acid-enhanced, multicellular human blood-brain barrier model derived from stem cell sources. Scientific Reports 4, 4160, doi:10.1038/srep04160 (2014).

40 Hollmann, E. K., Bailey, A. K., Potharazu, A. V., Neely, M. D., Bowman, A. B. & Lippmann, E. S. Accelerated differentiation of human induced pluripotent stem cells to blood–brain barrier endothelial cells. Fluids and Barriers of the CNS 14, 9, doi:10.1186/s12987-017-0059-0 (2017).

41 Qian, T., Maguire, S. E., Canfield, S. G., Bao, X., Olson, W. R., Shusta, E. V. & Palecek, S. P. Directed differentiation of human pluripotent stem cells to blood-brain barrier endothelial cells. Science Advances 3, e1701679, doi:10.1126/sciadv.1701679 (2017).

42 Linville, R. M., Nerenberg, R. F., Grifno, G., Arevalo, D., Guo, Z. & Searson, P. C. Brain microvascular endothelial cell dysfunction in an isogenic juvenile iPSC model of Huntington’s disease. Fluids and Barriers of the CNS 19, 54, doi:10.1186/s12987-022-00347-7 (2022).

43 Canfield, S. G., Stebbins, M. J., Morales, B. S., Asai, S. W., Vatine, G. D., Svendsen, C. N., Palecek, S. P. & Shusta, V. E. An isogenic blood–brain barrier model comprising brain endothelial cells, astrocytes, and neurons derived from human induced pluripotent stem cells. Journal of Neurochemistry 140, 874–888, doi:10.1111/jnc.13923 (2017).

44 Faley, S. L., Neal, E. H., Wang, J. X., Bosworth, A. M., Weber, C. M., Balotin, K. M., Lippmann, E. S. & Bellan, L. M. iPSC-Derived Brain Endothelium Exhibits Stable, Long-Term Barrier Function in Perfused Hydrogel Scaffolds. Stem cell reports 12, 474–487, doi:10.1016/j.stemcr.2019.01.009 (2019).

45 Kim, B. J., Bee, O. B., McDonagh, M. A., Stebbins, M. J., Palecek, S. P., Doran, K. S. & Shusta, E. V. Modeling Group B Streptococcus and Blood-Brain Barrier Interaction by Using Induced Pluripotent Stem Cell-Derived Brain Endothelial Cells. mSphere 2, e00398–00317, doi:10.1128/mSphere.00398-17 (2017).

46 Lim, R. G., Quan, C., Reyes-Ortiz, A. M., Lutz, S. E., Kedaigle, A. J., Gipson, T. A., Wu, J., Vatine, G. D., Stocksdale, J., Casale, M. S., Svendsen, C. N., Fraenkel, E., Housman, D. E., Agalliu, D. & Thompson, L. M. Huntington’s Disease iPSC-Derived Brain Microvascular Endothelial Cells Reveal WNT-Mediated Angiogenic and Blood-Brain Barrier Deficits. Cell reports 19, 1365–1377, doi:10.1016/j.celrep.2017.04.021 (2017).

47 Linville, R. M., DeStefano, J. G., Nerenberg, R. F., Grifno, G. N., Ye, R., Gallagher, E. & Searson, P. C. Long-Term Cryopreservation Preserves Blood–Brain Barrier Phenotype of iPSC-Derived Brain Microvascular Endothelial Cells and Three-Dimensional Microvessels. Molecular Pharmaceutics 17, 3425–3434, doi:10.1021/acs.molpharmaceut.0c00484 (2020).

48 Vatine, G. D., Al-Ahmad, A., Barriga, B. K., Svendsen, S., Salim, A., Garcia, L., Garcia, V. J., Ho, R., Yucer, N., Qian, T., Lim, R. G., Wu, J., Thompson, L. M., Spivia, W. R., Chen, Z., Van Eyk, J., Palecek, S. P., Refetoff, S., Shusta, E. V. & Svendsen, C. N. Modeling Psychomotor Retardation using iPSCs from MCT8-Deficient Patients Indicates a Prominent Role for the Blood-Brain Barrier. Cell Stem Cell 20, 831–843.e835, doi:10.1016/j.stem.2017.04.002 (2017).

49 Vatine, G. D., Barrile, R., Workman, M. J., Sances, S., Barriga, B. K., Rahnama, M., Barthakur, S., Kasendra, M., Lucchesi, C., Kerns, J., Wen, N., Spivia, W. R., Chen, Z., Van Eyk, J. & Svendsen, C. N. Human iPSC-Derived Blood-Brain Barrier Chips Enable Disease Modeling and Personalized Medicine Applications. Cell Stem Cell 24, 995–1005 e1006, doi:10.1016/j.stem.2019.05.011 (2019).

50 Pediaditakis, I., Kodella, K. R., Manatakis, D. V., Le, C. Y., Hinojosa, C. D., Tien-Street, W., Manolakos, E. S., Vekrellis, K., Hamilton, G. A., Ewart, L., Rubin, L. L. & Karalis, K. Modeling alpha-synuclein pathology in a human brain-chip to assess blood-brain barrier disruption. Nature Communications 12, 5907, doi:10.1038/s41467-021-26066-5 (2021).

51 Stebbins, M. J., Wilson, H. K., Canfield, S. G., Qian, T., Palecek, S. P. & Shusta, E. V. Differentiation and Characterization of Human Pluripotent Stem Cell-Derived Brain Microvascular Endothelial Cells. *Methods (San Diego*, Calif.) 101, 93–102, doi:10.1016/j.ymeth.2015.10.016 (2016).

52 Appelt-Menzel, A., Cubukova, A., Günther, K., Edenhofer, F., Piontek, J., Krause, G., Stüber, T., Walles, H., Neuhaus, W. & Metzger, M. Establishment of a Human Blood-Brain Barrier Co-culture Model Mimicking the Neurovascular Unit Using Induced Pluri- and Multipotent Stem Cells. Stem Cell Reports 8, 894–906, doi:10.1016/j.stemcr.2017.02.021 (2017).

53 Canfield, S. G., Stebbins, M. J., Morales, B. S., Asai, S. W., Vatine, G. D., Svendsen, C. N., Palecek, S. P. & Shusta, E. V. An Isogenic Blood-Brain Barrier Model Comprising Brain Endothelial Cells, Astrocytes and Neurons Derived from Human Induced Pluripotent Stem Cells. Journal of neurochemistry 140, 874–888, doi:10.1111/jnc.13923 (2017).

54 Stebbins Matthew, J., Lippmann Ethan, S., Faubion Madeline, G., Daneman, R., Palecek Sean, P. & Shusta Eric, V. Activation of RARα, RARγ, or RXRα Increases Barrier Tightness in Human Induced Pluripotent Stem Cell-Derived Brain Endothelial Cells. Biotechnology Journal 13, 1700093, doi:10.1002/biot.201700093 (2017).

55 Delsing, L., Donnes, P., Sanchez, J., Clausen, M., Voulgaris, D., Falk, A., Herland, A., Brolen, G., Zetterberg, H., Hicks, R. & Synnergren, J. Barrier Properties and Transcriptome Expression in Human iPSC-Derived Models of the Blood-Brain Barrier. Stem Cells 36, 1816–1827, doi:10.1002/stem.2908 (2018).

56 Lee, C. A. A., Seo, H. S., Armien, A. G., Bates, F. S., Tolar, J. & Azarin, S. M. Modeling and rescue of defective blood-brain barrier function of induced brain microvascular endothelial cells from childhood cerebral adrenoleukodystrophy patients. Fluids Barriers CNS 15, 9, doi:10.1186/s12987-018-0094-5 (2018).

57 Sances, S., Ho, R., Vatine, G., West, D., Laperle, A., Meyer, A., Godoy, M., Kay, P. S., Mandefro, B., Hatata, S., Hinojosa, C., Wen, N., Sareen, D., Hamilton, G. A. & Svendsen, C. N. Human iPSC-Derived Endothelial Cells and Microengineered Organ-Chip Enhance Neuronal Development. Stem Cell Reports 10, 1222–1236, doi:10.1016/j.stemcr.2018.02.012 (2018).

58 Martins Gomes, S. F., Westermann, A. J., Sauerwein, T., Hertlein, T., Forstner, K. U., Ohlsen, K., Metzger, M., Shusta, E. V., Kim, B. J., Appelt-Menzel, A. & Schubert-Unkmeir, A. Induced Pluripotent Stem Cell-Derived Brain Endothelial Cells as a Cellular Model to Study Neisseria meningitidis Infection. Front Microbiol 10, 1181, doi:10.3389/fmicb.2019.01181 (2019).

59 Wilson, H. K., Canfield, S. G., Hjortness, M. K., Palecek, S. P. & Shusta, E. V. Exploring the effects of cell seeding density on the differentiation of human pluripotent stem cells to brain microvascular endothelial cells. Fluids and Barriers of the CNS 12, 13, doi:10.1186/s12987-015-0007-9 (2015).

60 Lippmann, E. S., Al-Ahmad, A., Palecek, S. P. & Shusta, E. V. Modeling the blood–brain barrier using stem cell sources. Fluids and Barriers of the CNS 10, 2–2, doi:10.1186/2045-8118-10-2 (2013).

61 Abbott, N. J. Astrocyte–endothelial interactions and blood–brain barrier permeability. Journal of Anatomy 200, 629–638, 10.1046/j.1469-7580.2002.00064.x (2002).

62 Siletti, K., Hodge, R., Mossi Albiach, A., Lee, K. W., Ding, S. L., Hu, L., Lönnerberg, P., Bakken, T., Casper, T., Clark, M., Dee, N., Gloe, J., Hirschstein, D., Shapovalova, N. V., Keene, C. D., Nyhus, J., Tung, H., Yanny, A. M., Arenas, E., Lein, E. S. & Linnarsson, S. Transcriptomic diversity of cell types across the adult human brain. Science 382, eadd7046, doi:10.1126/science.add7046 (2023).

63 Yang, A. C., Vest, R. T., Kern, F., Lee, D. P., Agam, M., Maat, C. A., Losada, P. M., Chen, M. B., Schaum, N., Khoury, N., Toland, A., Calcuttawala, K., Shin, H., Pálovics, R., Shin, A., Wang, E. Y., Luo, J., Gate, D., Schulz-Schaeffer, W. J., Chu, P., Siegenthaler, J. A., McNerney, M. W., Keller, A. & Wyss-Coray, T. A human brain vascular atlas reveals diverse mediators of Alzheimer’s risk. Nature 603, 885–892, doi:10.1038/s41586-021-04369-3 (2022).

64 Lin, Y., Gil, C.-H. & Yoder, M. C. Differentiation, Evaluation, and Application of Human Induced Pluripotent Stem Cell–Derived Endothelial Cells. Arteriosclerosis, Thrombosis, and Vascular Biology 37, 2014–2025, doi:10.1161/ATVBAHA.117.309962 (2017).

65 Oh, S. Y., Kim, J. Y. & Park, C. The ETS factor, ETV2: A master regulator for vascular endothelial cell development. Molecules and Cells 38, 1029–1036, doi:10.14348/molcells.2015.0331 (2015).

66 Palikuqi, B., Nguyen, D. T., Li, G., Schreiner, R., Pellegata, A. F., Liu, Y., Redmond, D., Geng, F., Lin, Y., Gómez-Salinero, J. M., Yokoyama, M., Zumbo, P., Zhang, T., Kunar, B., Witherspoon, M., Han, T., Tedeschi, A. M., Scottoni, F., Lipkin, S. M., Dow, L., Elemento, O., Xiang, J. Z., Shido, K., Spence, J. R., Zhou, Q. J., Schwartz, R. E., De Coppi, P., Rabbany, S. Y. & Rafii, S. Adaptable haemodynamic endothelial cells for organogenesis and tumorigenesis. Nature 585, 426–432, doi:10.1038/s41586-020-2712-z (2020).

67 Kataoka, H., Hayashi, M., Nakagawa, R., Tanaka, Y., Izumi, N., Nishikawa, S., Jakt, M. L., Tarui, H. & Nishikawa, S.-I. Etv2/ER71 induces vascular mesoderm from Flk1+PDGFRα+ primitive mesoderm. Blood 118, 6975–6986, 10.1182/blood-2011-05-352658 (2011).

68 Liu, F., Li, D., Yu, Y. Y. L., Kang, I., Cha, M. J., Kim, J. Y., Park, C., Watson, D. K., Wang, T. & Choi, K. Induction of hematopoietic and endothelial cell program orchestrated by ETS transcription factor ER71/ETV2. EMBO reports 16, 654–669–669, 10.15252/embr.201439939 (2015).

69 Ferdous, A., Caprioli, A., Iacovino, M., Martin, C. M., Morris, J., Richardson, J. A., Latif, S., Hammer, R. E., Harvey, R. P., Olson, E. N., Kyba, M. & Garry, D. J. Nkx2–5 transactivates the Ets-related protein 71 gene and specifies an endothelial/endocardial fate in the developing embryo. Proceedings of the National Academy of Sciences 106, 814–819, doi:10.1073/pnas.0807583106 (2009).

70 Lee, D., Park, C., Lee, H., Lugus, J. J., Kim, S. H., Arentson, E., Chung, Y. S., Gomez, G., Kyba, M., Lin, S., Janknecht, R., Lim, D.-S. & Choi, K. ER71 Acts Downstream of BMP, Notch, and Wnt Signaling in Blood and Vessel Progenitor Specification. Cell Stem Cell 2, 497–507, doi:10.1016/j.stem.2008.03.008 (2008).

71 Oh, S.-Y., Kim, J. Y. & Park, C. The ETS Factor, ETV2: a Master Regulator for Vascular Endothelial Cell Development. Molecules and Cells 38, 1029–1036, 10.14348/molcells.2015.0331 (2015).

72 Luo, A. C., Wang, J., Wang, K., Zhu, Y., Gong, L., Lee, U., Li, X., Tremmel, D. M., Lin, R. Z., Ingber, D. E., Gorman, J. & Melero-Martin, J. M. A streamlined method to generate endothelial cells from human pluripotent stem cells via transient doxycycline-inducible ETV2 activation. Angiogenesis 27, 779–795, doi:10.1007/s10456-024-09937-5 (2024).

73 Wang, K., Lin, R. Z., Hong, X., Ng, A. H., Lee, C. N., Neumeyer, J., Wang, G., Wang, X., Ma, M., Pu, W. T., Church, G. M. & Melero-Martin, J. M. Robust differentiation of human pluripotent stem cells into endothelial cells via temporal modulation of ETV2 with modified mRNA. Sci Adv 6, eaba7606, doi:10.1126/sciadv.aba7606 (2020).

74 Birdsey, Graeme M., Shah, Aarti V., Dufton, N., Reynolds, Louise E., Osuna Almagro, L., Yang, Y., Aspalter, Irene M., Khan, Samia T., Mason, Justin C., Dejana, E., Göttgens, B., Hodivala-Dilke, K., Gerhardt, H., Adams, Ralf H. & Randi, Anna M. The Endothelial Transcription Factor ERG Promotes Vascular Stability and Growth through Wnt/&#x3b2;-Catenin Signaling. Developmental Cell 32, 82–96, doi:10.1016/j.devcel.2014.11.016 (2015).

75 Kalna, V., Yang, Y., Peghaire, C. R., Frudd, K., Hannah, R., Shah, A. V., Osuna Almagro, L., Boyle, J. J., Göttgens, B., Ferrer, J., Randi, A. M. & Birdsey, G. M. The Transcription Factor ERG Regulates Super-Enhancers Associated With an Endothelial-Specific Gene Expression Program. Circulation Research 124, 1337–1349, doi:10.1161/CIRCRESAHA.118.313788 (2019).

76 Shah, A. V., Birdsey, G. M., Peghaire, C., Pitulescu, M. E., Dufton, N. P., Yang, Y., Weinberg, I., Osuna Almagro, L., Payne, L., Mason, J. C., Gerhardt, H., Adams, R. H. & Randi, A. M. The endothelial transcription factor ERG mediates Angiopoietin-1-dependent control of Notch signalling and vascular stability. Nature Communications 8, 16002, doi:10.1038/ncomms16002 (2017).

77 Shah, A. V., Birdsey, G. M. & Randi, A. M. Regulation of endothelial homeostasis, vascular development and angiogenesis by the transcription factor ERG. Vascular Pharmacology 86, 3– 13, 10.1016/j.vph.2016.05.003 (2016).

78 Ginsberg, M., James, D., Ding, S. B., Nolan, D., Geng, F., Butler, J. M., Schachterle, W., Pulijaal, V. R., Mathew, S., Chasen, S. T., Xiang, J., Rosenwaks, Z., Shido, K., Elemento, O., Rabbany, S. Y. & Rafii, S. Efficient direct reprogramming of mature amniotic cells into endothelial cells by ETS factors and TGFβ suppression. Cell 151, 559–575, doi:10.1016/j.cell.2012.09.032 (2012).

79 Ginsberg, M., Schachterle, W., Shido, K. & Rafii, S. Direct conversion of human amniotic cells into endothelial cells without transitioning through a pluripotent state. Nature Protocols 10, 1975, doi:10.1038/nprot.2015.126 https://www.nature.com/articles/nprot.2015.126#supplementary-information (2015).

80 Lu, T. M., Houghton, S., Magdeldin, T., Durán, J. G. B., Minotti, A. P., Snead, A., Sproul, A., Nguyen, D.-H. T., Xiang, J., Fine, H. A., Rosenwaks, Z., Studer, L., Rafii, S., Agalliu, D., Redmond, D. & Lis, R. Pluripotent stem cell-derived epithelium misidentified as brain microvascular endothelium requires ETS factors to acquire vascular fate. Proceedings of the National Academy of Sciences 118, e2016950118, doi:10.1073/pnas.2016950118 (2021).

81 Sarkar, A. & Hochedlinger, K. The sox family of transcription factors: versatile regulators of stem and progenitor cell fate. Cell Stem Cell 12, 15–30, doi:10.1016/j.stem.2012.12.007 (2013).

82 De Val, S. & Black, B. L. Transcriptional Control of Endothelial Cell Development. Developmental Cell 16, 180–195, doi:10.1016/j.devcel.2009.01.014 (2009).

83 Guth, S. I. & Wegner, M. Having it both ways: Sox protein function between conservation and innovation. Cell Mol Life Sci 65, 3000–3018, doi:10.1007/s00018-008-8138-7 (2008).

84 Kanki, Y., Nakaki, R., Shimamura, T., Matsunaga, T., Yamamizu, K., Katayama, S., Suehiro, J.-i., Osawa, T., Aburatani, H., Kodama, T., Wada, Y., Yamashita, J. K. & Minami, T. Dynamically and epigenetically coordinated GATA/ETS/SOX transcription factor expression is indispensable for endothelial cell differentiation. Nucleic Acids Research 45, 4344–4358, doi:10.1093/nar/gkx159 (2017).

85 Chiang, I. K.-N., Fritzsche, M., Pichol-Thievend, C., Neal, A., Holmes, K., Lagendijk, A., Overman, J., D’Angelo, D., Omini, A., Hermkens, D., Lesieur, E., Liu, K., Ratnayaka, I., Corada, M., Bou-Gharios, G., Carroll, J., Dejana, E., Schulte-Merker, S., Hogan, B., Beltrame, M., De Val, S. & Francois, M. SoxF factors induce Notch1 expression via direct transcriptional regulation during early arterial development. Development 144, 2629–2639, doi:10.1242/dev.146241 (2017).

86 Francois, M., Koopman, P. & Beltrame, M. SoxF genes: Key players in the development of the cardio-vascular system. The International Journal of Biochemistry & Cell Biology 42, 445–448, 10.1016/j.biocel.2009.08.017 (2010).

87 Kim, K., Kim, I.-K., Yang, J. M., Lee, E., Koh, B. I., Song, S., Park, J., Lee, S., Choi, C., Kim, J. W., Kubota, Y., Koh, G. Y. & Kim, I. SoxF Transcription Factors Are Positive Feedback Regulators of VEGF Signaling. Circulation Research 119, 839–852, doi:10.1161/CIRCRESAHA.116.308483 (2016).

88 Patel, J., Seppanen, E. J., Rodero, M. P., Wong, H. Y., Donovan, P., Neufeld, Z., Fisk, N. M., Francois, M. & Khosrotehrani, K. Functional Definition of Progenitors Versus Mature Endothelial Cells Reveals Key SoxF-Dependent Differentiation Process. Circulation 135, 786–805, doi:10.1161/CIRCULATIONAHA.116.024754 (2017).

89 Zhou, Y., Williams, J., Smallwood, P. M. & Nathans, J. Sox7, Sox17, and Sox18 Cooperatively Regulate Vascular Development in the Mouse Retina. PLOS ONE 10, e0143650, doi:10.1371/journal.pone.0143650 (2015).

90 Farber, G., Takasugi, P., Ricketts, S., Wang, H., Xie, Y., Farber, E., Liu, J. & Qian, L. Sox17 and Erg synergistically activate endothelial cell fate in reprogramming fibroblasts. Journal of Molecular and Cellular Cardiology 199, 33–45, 10.1016/j.yjmcc.2024.11.012 (2025).

91 Schaffenrath, J., Wyss, T., He, L., Rushing, E. J., Delorenzi, M., Vasella, F., Regli, L., Neidert, M. C. & Keller, A. Blood-brain barrier alterations in human brain tumors revealed by genome-wide transcriptomic profiling. Neuro Oncol 23, 2095–2106, doi:10.1093/neuonc/noab022 (2021).

92 Zhang, Y., Sloan, S. A., Clarke, L. E., Caneda, C., Plaza, C. A., Blumenthal, P. D., Vogel, H., Steinberg, G. K., Edwards, M. S., Li, G., Duncan, J. A., 3rd, Cheshier, S. H., Shuer, L. M., Chang, E. F., Grant, G. A., Gephart, M. G. & Barres, B. A. Purification and Characterization of Progenitor and Mature Human Astrocytes Reveals Transcriptional and Functional Differences with Mouse. Neuron 89, 37–53, doi:10.1016/j.neuron.2015.11.013 (2016).

93 Elizarraras, J. M., Liao, Y., Shi, Z., Zhu, Q., Pico, Alexander R. & Zhang, B. WebGestalt 2024: faster gene set analysis and new support for metabolomics and multi-omics. Nucleic Acids Research 52, W415–W421, doi:10.1093/nar/gkae456 (2024).

94 Zhang, B., Kirov, S. & Snoddy, J. WebGestalt: an integrated system for exploring gene sets in various biological contexts. Nucleic Acids Research 33, W741–W748, doi:10.1093/nar/gki475 (2005).

95 Leucht, P., Kim, J.-B., Amasha, R., James, A. W., Girod, S. & Helms, J. A. Embryonic origin and Hox status determine progenitor cell fate during adult bone regeneration. Development 135, 2845–2854, doi:10.1242/dev.023788 (2008).

96 Lippmann, E. S., Estevez-Silva, M. C. & Ashton, R. S. Defined Human Pluripotent Stem Cell Culture Enables Highly Efficient Neuroepithelium Derivation Without Small Molecule Inhibitors. Stem Cells 32, 1032–1042, doi:10.1002/stem.1622 (2014).

97 de Sousa Abreu, R., Penalva, L. O., Marcotte, E. M. & Vogel, C. Global signatures of protein and mRNA expression levels. Mol Biosyst 5, 1512–1526, doi:10.1039/b908315d (2009).

98 Maier, T., Güell, M. & Serrano, L. Correlation of mRNA and protein in complex biological samples. FEBS Letters 583, 3966–3973, 10.1016/j.febslet.2009.10.036 (2009).

99 Vogel, C. & Marcotte, E. M. Insights into the regulation of protein abundance from proteomic and transcriptomic analyses. Nature Reviews Genetics 13, 227–232, doi:10.1038/nrg3185 (2012).

100 Bauernfeind, A. L. & Babbitt, C. C. The predictive nature of transcript expression levels on protein expression in adult human brain. BMC Genomics 18, 322, doi:10.1186/s12864-017-3674-x (2017).

101 Ben-Zvi, A., Lacoste, B., Kur, E., Andreone, B. J., Mayshar, Y., Yan, H. & Gu, C. Mfsd2a is critical for the formation and function of the blood-brain barrier. Nature 509, 507–511, doi:10.1038/nature13324 (2014).

102 Daneman, R., Zhou, L., Kebede, A. A. & Barres, B. A. Pericytes are required for blood-brain barrier integrity during embryogenesis. Nature 468, 562–566, doi:10.1038/nature09513 (2010).

103 Creuzet, S., Couly, G., Vincent, C. & Le Douarin, N. M. Negative effect of Hox gene expression on the development of the neural crest-derived facial skeleton. Development 129, 4301–4313, doi:10.1242/dev.129.18.4301 (2002).

104 Le Douarin, N. M., Creuzet, S., Couly, G. & Dupin, E. Neural crest cell plasticity and its limits. Development 131, 4637–4650, doi:10.1242/dev.01350 (2004).

105 Nolan, D. J., Ginsberg, M., Israely, E., Palikuqi, B., Poulos, M. G., James, D., Ding, B.-S., Schachterle, W., Liu, Y., Rosenwaks, Z., Butler, J. M., Xiang, J., Rafii, A., Shido, K., Rabbany, S. Y., Elemento, O. & Rafii, S. Molecular signatures of tissue-specific microvascular endothelial cell heterogeneity in organ maintenance and regeneration. Developmental Cell 26, 204–219, doi:10.1016/j.devcel.2013.06.017 (2013).

106 Toshner, M., Dunmore, B. J., McKinney, E. F., Southwood, M., Caruso, P., Upton, P. D., Waters, J. P., Ormiston, M. L., Skepper, J. N., Nash, G., Rana, A. A. & Morrell, N. W. Transcript Analysis Reveals a Specific HOX Signature Associated with Positional Identity of Human Endothelial Cells. PLOS ONE 9, e91334, doi:10.1371/journal.pone.0091334 (2014).

107 Costa, G., Mazan, A., Gandillet, A., Pearson, S., Lacaud, G. & Kouskoff, V. SOX7 regulates the expression of VE-cadherin in the haemogenic endothelium at the onset of haematopoietic development. Development 139, 1587–1598, doi:10.1242/dev.071282 (2012).

108 Munji, R. N., Soung, A. L., Weiner, G. A., Sohet, F., Semple, B. D., Trivedi, A., Gimlin, K., Kotoda, M., Korai, M., Aydin, S., Batugal, A., Cabangcala, A. C., Schupp, P. G., Oldham, M. C., Hashimoto, T., Noble-Haeusslein, L. J. & Daneman, R. Profiling the mouse brain endothelial transcriptome in health and disease models reveals a core blood-brain barrier dysfunction module. Nat Neurosci 22, 1892–1902, doi:10.1038/s41593-019-0497-x (2019).

109 Rafii, S., Shapiro, F., Pettengell, R., Ferris, B., Nachman, R. L., Moore, M. A. & Asch, A. S. Human bone marrow microvascular endothelial cells support long-term proliferation and differentiation of myeloid and megakaryocytic progenitors. Blood 86, 3353 (1995).

110 Butler, J. M. & Rafii, S. Generation of a vascular niche for studying stem cell homeostasis. Methods Mol Biol 904, 221–233, doi:10.1007/978-1-61779-943-3_18 (2012).

111 Yu, J., Vodyanik, M. A., Smuga-Otto, K., Antosiewicz-Bourget, J., Frane, J. L., Tian, S., Nie, J., Jonsdottir, G. A., Ruotti, V., Stewart, R., Slukvin, I. I. & Thomson, J. A. Induced Pluripotent Stem Cell Lines Derived from Human Somatic Cells. Science 318, 1917, doi:10.1126/science.1151526 (2007).

112 Yu, J., Hu, K., Smuga-Otto, K., Tian, S., Stewart, R., Slukvin, I. I. & Thomson, J. A. Human Induced Pluripotent Stem Cells Free of Vector and Transgene Sequences. Science 324, 797, doi:10.1126/science.1172482 (2009).

113 Chen, G., Gulbranson, D. R., Hou, Z., Bolin, J. M., Ruotti, V., Probasco, M. D., Smuga-Otto, K., Howden, S. E., Diol, N. R., Propson, N. E., Wagner, R., Lee, G. O., Antosiewicz-Bourget, J., Teng, J. M. C. & Thomson, J. A. Chemically defined conditions for human iPSC derivation and culture. Nature Methods 8, 424–429, doi:10.1038/nmeth.1593 (2011).

114 Hu, K., Yu, J., Suknuntha, K., Tian, S., Montgomery, K., Choi, K.-D., Stewart, R., Thomson, J. A. & Slukvin, I. I. Efficient generation of transgene-free induced pluripotent stem cells from normal and neoplastic bone marrow and cord blood mononuclear cells. Blood 117, e109–e119, doi:10.1182/blood-2010-07-298331 (2011).

115 Ziemann, M., Kaspi, A. & El-Osta, A. Digital expression explorer 2: a repository of uniformly processed RNA sequencing data. Gigascience 8, giz022, doi:10.1093/gigascience/giz022 (2019).

116 Jiang, H., Lei, R., Ding, S.-W. & Zhu, S. Skewer: a fast and accurate adapter trimmer for next-generation sequencing paired-end reads. BMC Bioinformatics 15, 182–182, doi:10.1186/1471-2105-15-182 (2014).

117 Davis, M. P., van Dongen, S., Abreu-Goodger, C., Bartonicek, N. & Enright, A. J. Kraken: a set of tools for quality control and analysis of high-throughput sequence data. Methods 63, 41–49, doi:10.1016/j.ymeth.2013.06.027 (2013).

118 Dobin, A., Davis, C. A., Schlesinger, F., Drenkow, J., Zaleski, C., Jha, S., Batut, P., Chaisson, M. & Gingeras, T. R. STAR: ultrafast universal RNA-seq aligner. Bioinformatics 29, 15–21, doi:10.1093/bioinformatics/bts635 (2013).

119 Robinson, M. D., McCarthy, D. J. & Smyth, G. K. edgeR: a Bioconductor package for differential expression analysis of digital gene expression data. *Bioinformatics (Oxford*, England) 26, 139– 140, doi:10.1093/bioinformatics/btp616 (2010).

120 Ritchie, M. E., Phipson, B., Wu, D., Hu, Y., Law, C. W., Shi, W. & Smyth, G. K. limma powers differential expression analyses for RNA-sequencing and microarray studies. Nucleic Acids Research 43, e47–e47, doi:10.1093/nar/gkv007 (2015).

121 Liao, Y., Wang, J., Jaehnig, E. J., Shi, Z. & Zhang, B. WebGestalt 2019: gene set analysis toolkit with revamped UIs and APIs. Nucleic acids research 47, W199–W205, doi:10.1093/nar/gkz401 (2019).

122 Li, J., Cai, Z., Bomgarden, R. D., Pike, I., Kuhn, K., Rogers, J. C., Roberts, T. M., Gygi, S. P. & Paulo, J. A. TMTpro-18plex: The Expanded and Complete Set of TMTpro Reagents for Sample Multiplexing. J Proteome Res 20, 2964–2972, doi:10.1021/acs.jproteome.1c00168 (2021).

123 Navarrete-Perea, J., Yu, Q., Gygi, S. P. & Paulo, J. A. Streamlined Tandem Mass Tag (SL-TMT) Protocol: An Efficient Strategy for Quantitative (Phospho)proteome Profiling Using Tandem Mass Tag-Synchronous Precursor Selection-MS3. J Proteome Res 17, 2226–2236, doi:10.1021/acs.jproteome.8b00217 (2018).

124 Ginsberg, M., James, D., Ding, B.-S., Nolan, D., Geng, F., Butler, J. M., Schachterle, W., Pulijaal, V. R., Mathew, S., Chasen, S. T., Xiang, J., Rosenwaks, Z., Shido, K., Elemento, O., Rabbany, S. Y. & Rafii, S. Efficient direct reprogramming of mature amniotic cells into endothelial cells by ETS factors and TGFβ suppression. Cell 151, 559–575, doi:10.1016/j.cell.2012.09.032 (2012).

