## Supplemental Figures for "SOX7 and ERG reprograms neuroectoderm into microenvironmentally attuned brain endothelial cells that possess forebrain regional bias": FS1.pdf

### Figure S1

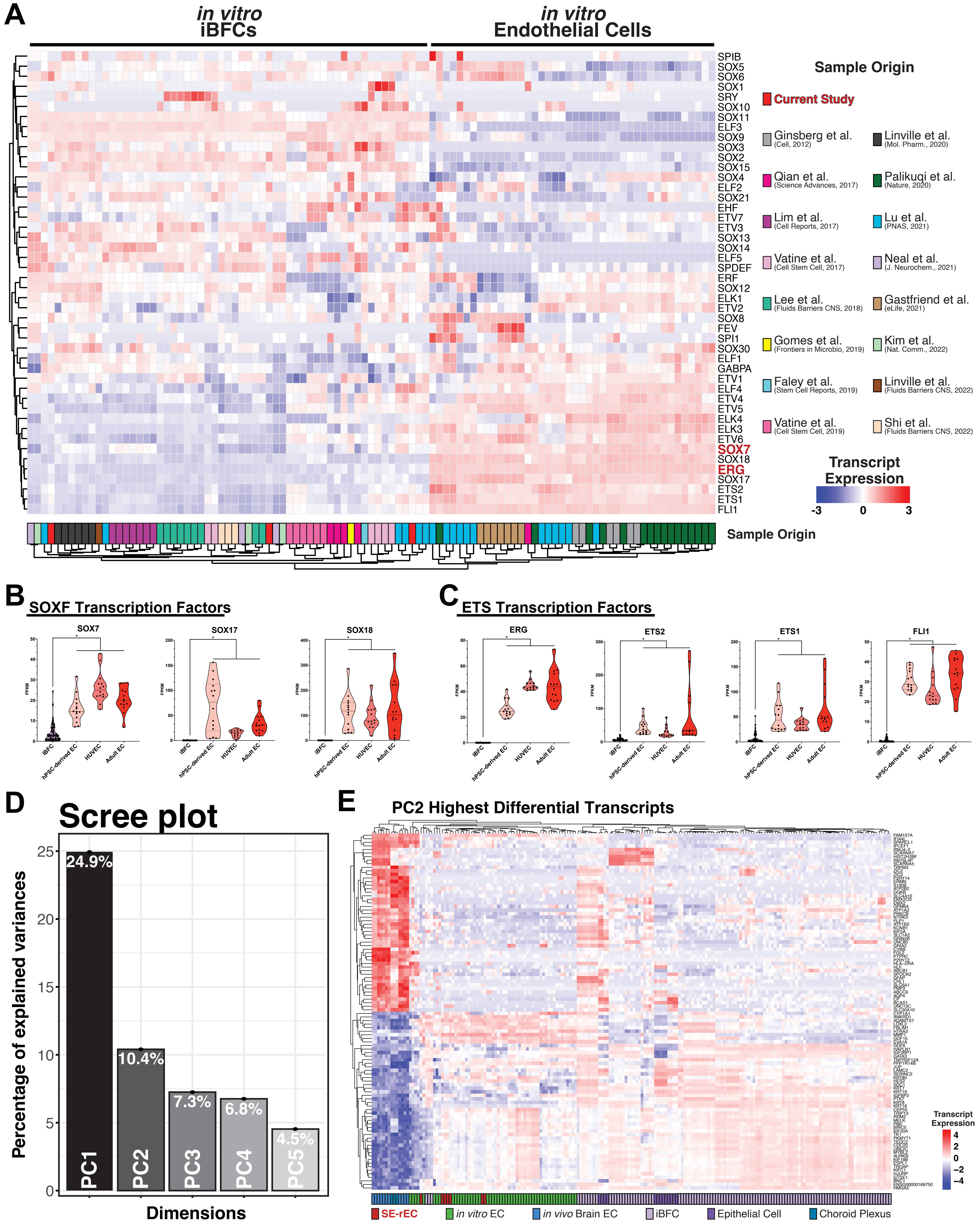
A
in vitro
iBFCs
in vitro
Endothelial Cells
SPIB
SOX5
SOX6
SOX1
SRY
SOX10
SOX11
ELF3
SOX9
SOX3
SOX2
SOX15
SOX4
ELF2
SOX21
EHF
ETV7
ETV3
SOX13
SOX14
ELF5
SPDEF
ERF
SOX12
ELK1
ETV2
SOX8
FEV
SPI1
SOX30
ELF1
GABPA
ETV1
ELF4
ETV4
ETV5
ELK4
ELK3
ETV6
SOX7
SOX18
ERG
SOX17
ETS2
ETS1
FLI1
Sample Origin
Current Study
Ginsberg et al. (Cell, 2012)
Linville et al. (Mol. Pharm., 2020)
Qian et al. (Science Advances, 2017)
Palikuqi et al. (Nature, 2020)
Lim et al. (Cell Reports, 2017)
Lu et al. (PNAS, 2021)
Vatine et al. (Cell Stem Cell, 2017)
Neal et al. (J. Neurochem., 2021)
Lee et al. (Fluids Barriers CNS, 2018)
Gastfriend et al. (eLife, 2021)
Gomes et al. (Frontiers in Microbio, 2019)
Kim et al. (Nat. Comm., 2022)
Faley et al. (Stem Cell Reports, 2019)
Linville et al. (Fluids Barriers CNS, 2022)
Vatine et al. (Cell Stem Cell, 2019)
Shi et al. (Fluids Barriers CNS, 2022)
Transcript Expression
-3
0
3
Sample Origin
B
SOXF Transcription Factors
SOX7
SOX17
SOX18
C
ETS Transcription Factors
ERG
ETS2
ETS1
FLI1
FPKM
iBFC
hPSC-derived EC
HUVEC
Adult EC
D
Scree plot
Percentage of explained variances
Dimensions
24.9%
10.4%
7.3%
6.8%
4.5%
PC1
PC2
PC3
PC4
PC5
E
PC2 Highest Differential Transcripts
Transcript Expression
4
2
0
−2
−4
SE-rEC
in vitro EC
in vivo Brain EC
iBFC
Epithelial Cell
Choroid Plexus
