## Supplemental Figures for "SOX7 and ERG reprograms neuroectoderm into microenvironmentally attuned brain endothelial cells that possess forebrain regional bias": FS3.pdf

### Figure S3

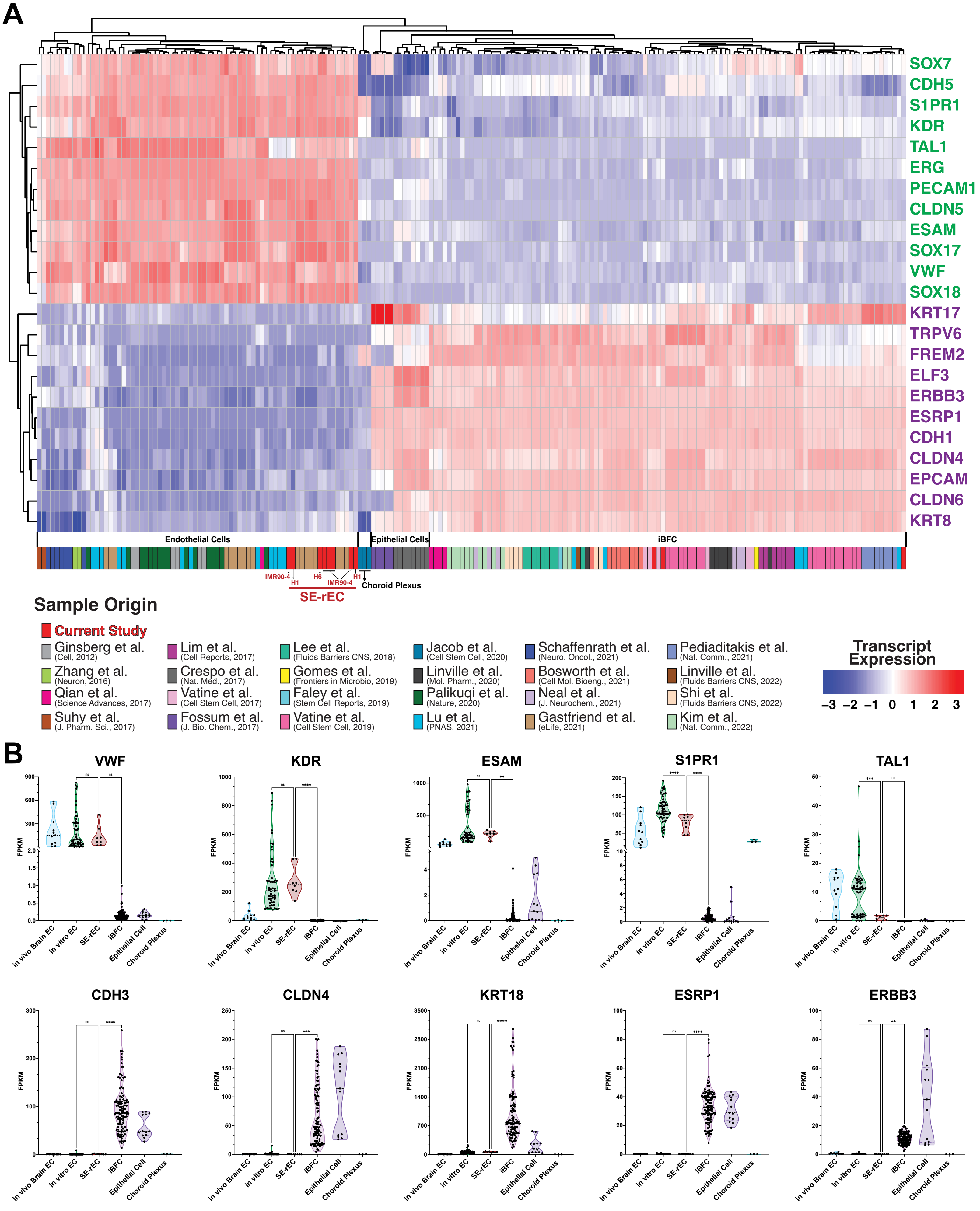
A
SOX7
CDH5
S1PR1
KDR
TAL1
ERG
PECAM1
CLDN5
ESAM
SOX17
VWF
SOX18
KRT17
TRPV6
FREM2
ELF3
ERBB3
ESRP1
CDH1
CLDN4
EPCAM
CLDN6
KRT8
Endothelial Cells
Epithelial Cells
iBFC
IMR90-4
H1
H6
IMR90-4
H1
Choroid Plexus
SE-rEC
Sample Origin
Current Study
Ginsberg et al. (Cell, 2012)
Zhang et al. (Neuron, 2016)
Qian et al. (Science Advances, 2017)
Suhy et al. (J. Pharm. Sci., 2017)
Lim et al. (Cell Reports, 2017)
Crespo et al. (Nat. Med., 2017)
Vatine et al. (Cell Stem Cell, 2017)
Fossum et al. (J. Bio. Chem., 2017)
Lee et al. (Fluids Barriers CNS, 2018)
Gomes et al. (Frontiers in Microbio, 2019)
Faley et al. (Stem Cell Reports, 2019)
Vatine et al. (Cell Stem Cell, 2019)
Jacob et al. (Cell Stem Cell, 2020)
Linville et al. (Mol. Pharm., 2020)
Palikuqi et al. (Nature, 2020)
Lu et al. (PNAS, 2021)
Schaffenrath et al. (Neuro. Oncol., 2021)
Bosworth et al. (Cell Mol. Bioeng., 2021)
Neal et al. (J. Neurochem., 2021)
Gastfriend et al. (eLife, 2021)
Pediaditakis et al. (Nat. Comm., 2021)
Linville et al. (Fluids Barriers CNS, 2022)
Shi et al. (Fluids Barriers CNS, 2022)
Kim et al. (Nat. Comm., 2022)
Transcript Expression
−3 −2 −1 0 1 2 3
B
VWF
KDR
ESAM
S1PR1
TAL1
CDH3
CLDN4
KRT18
ESRP1
ERBB3
FPKM
in vivo Brain EC
in vitro EC
SE-rEC
iBFC
Epithelial Cell
Choroid Plexus
ns
****
***
**
