## Supplemental Figures for "SOX7 and ERG reprograms neuroectoderm into microenvironmentally attuned brain endothelial cells that possess forebrain regional bias": FS4.pdf

### Figure S4

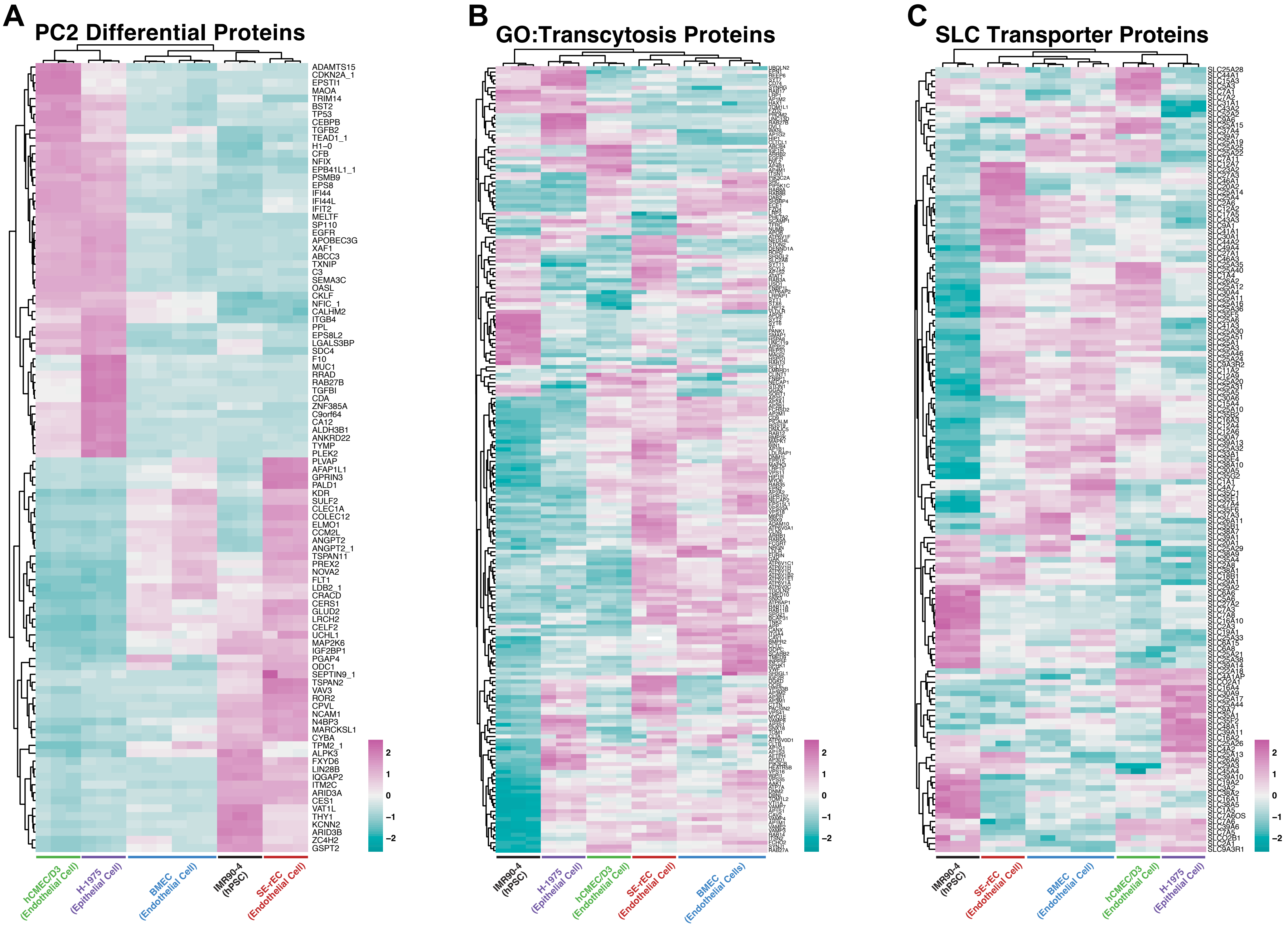

A
PC2 Differential Proteins
ADAMTS15
CDKN2A_1
EPSTI1
MAOA
TRIM14
BST2
TP53
CEBPB
TGFB2
TEAD1_1
H1−0
CFB
NFIX
EPB41L1_1
PSMB9
EPS8
IFI44
IFI44L
IFIT2
MELTF
SP110
EGFR
APOBEC3G
XAF1
ABCC3
TXNIP
C3
SEMA3C
OASL
CKLF
NFIC_1
CALHM2
ITGB4
PPL
EPS8L2
LGALS3BP
SDC4
F10
MUC1
RRAD
RAB27B
TGFBI
CDA
ZNF385A
C9orf64
CA12
ALDH3B1
ANKRD22
TYMP
PLEK2
PLVAP
AFAP1L1
GPRIN3
PALD1
KDR
SULF2
CLEC1A
COLEC12
ELMO1
CCM2L
ANGPT2
ANGPT2_1
TSPAN11
PREX2
NOVA2
FLT1
LDB2_1
CRACD
CERS1
GLUD2
LRCH2
CELF2
UCHL1
MAP2K6
IGF2BP1
PGAP4
ODC1
SEPTIN9_1
TSPAN2
VAV3
ROR2
CPVL
NCAM1
N4BP3
MARCKSL1
CYBA
TPM2_1
ALPK3
FXYD6
LIN28B
IQGAP2
ITM2C
ARID3A
CES1
VAT1L
THY1
KCNN2
ARID3B
ZC4H2
GSPT2
2
1
0
−1
−2
hCMEC/D3 (Endothelial Cell)
H-1975 (Epithelial Cell)
BMEC (Endothelial Cell)
IMR90-4 (hPSC)
SE-rEC (Endothelial Cell)
B
GO:Transcytosis Proteins
2
1
0
−1
−2
IMR90-4 (hPSC)
H-1975 (Epithelial Cell)
hCMEC/D3 (Endothelial Cell)
SE-rEC (Endothelial Cell)
BMEC (Endothelial Cells)
C
SLC Transporter Proteins
2
1
0
−1
−2
IMR90-4 (hPSC)
SE-rEC (Endothelial Cell)
BMEC (Endothelial Cell)
hCMEC/D3 (Endothelial Cell)
H-1975 (Epithelial Cell)
