## Supplemental Figures for "SOX7 and ERG reprograms neuroectoderm into microenvironmentally attuned brain endothelial cells that possess forebrain regional bias": FS5.pdf

### Figure S5

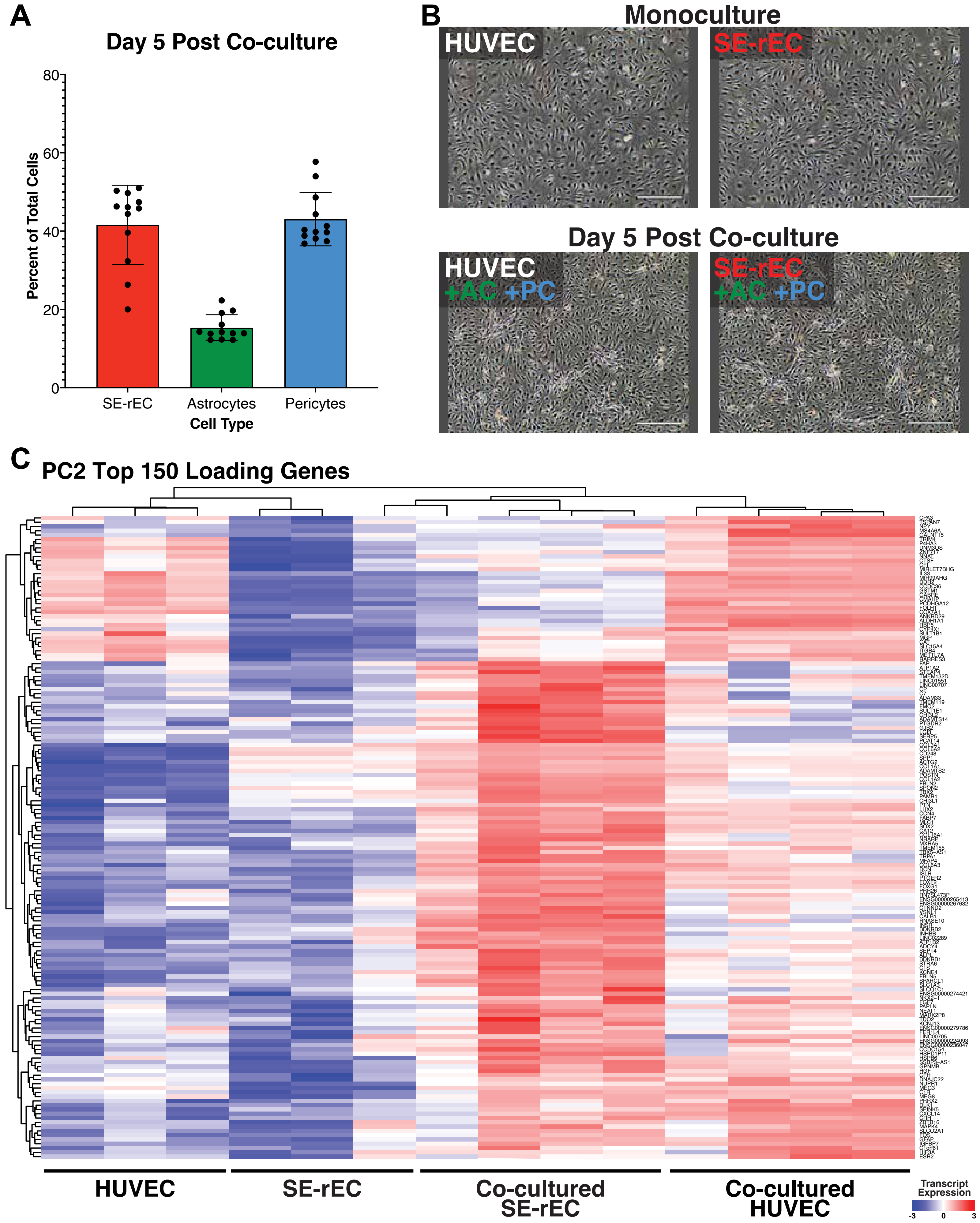
A
Day 5 Post Co-culture
Percent of Total Cells
80
60
40
20
0
SE-rEC
Astrocytes
Pericytes
Cell Type
B
Monoculture
HUVEC
SE-rEC
Day 5 Post Co-culture
HUVEC
+AC +PC
SE-rEC
+AC +PC
C
PC2 Top 150 Loading Genes
HUVEC
SE-rEC
Co-cultured SE-rEC
Co-cultured HUVEC
Transcript Expression
-3
0
3
