## Supplementary figures and images for "SOX7 and ERG reprograms neuroectoderm into microenvironmentally attuned brain endothelial cells that possess forebrain regional bias"

### FS2.pdf

#### Figure S2

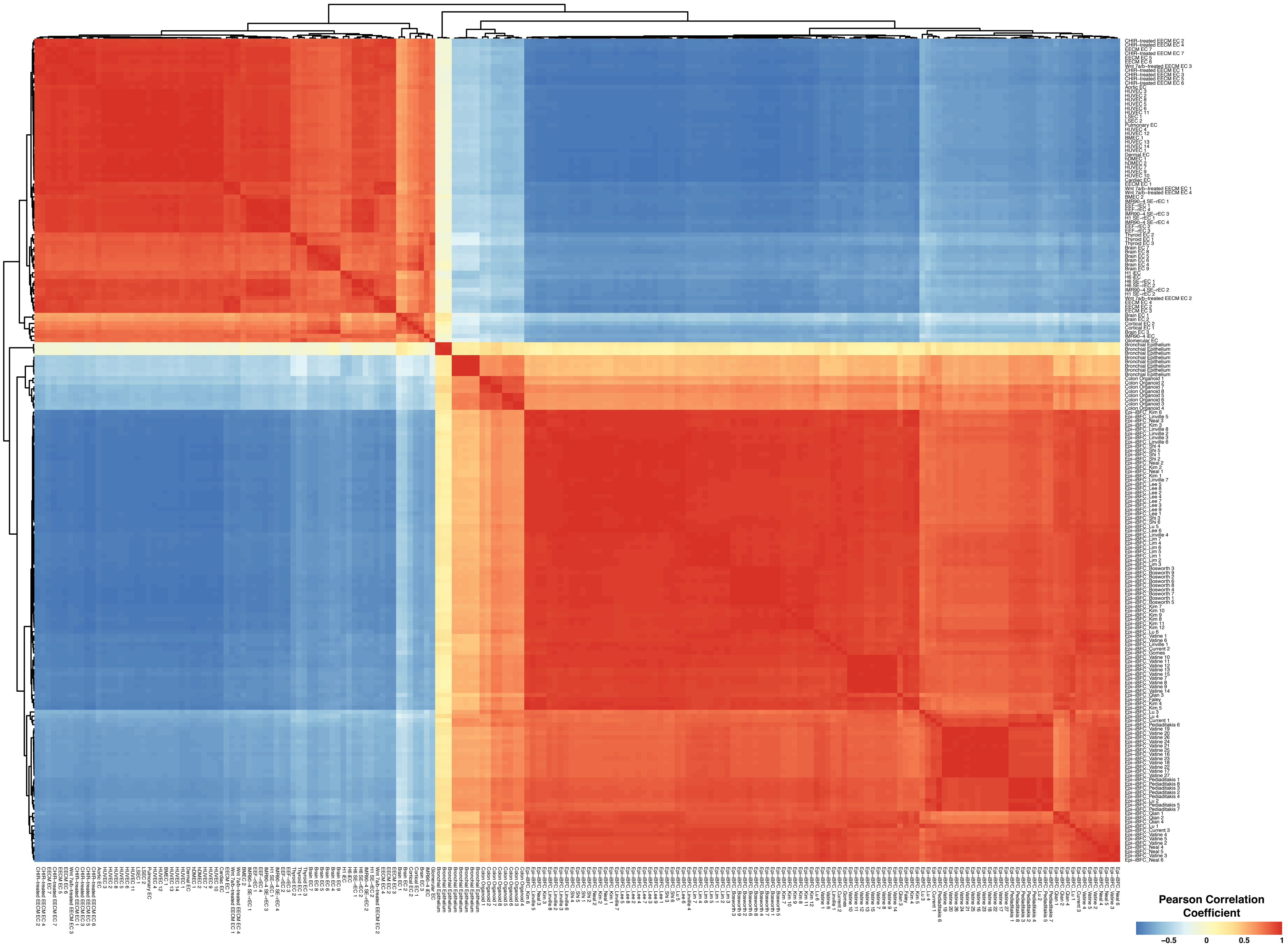
